# Frequency modulation of organelle calcium-dependent voltage oscillations provides an essential memory trace of operant conditioning in *Aplysia*

**DOI:** 10.64898/2026.07.30.741715

**Authors:** Laura Puygrenier, John Simmers, Romuald Nargeot

## Abstract

Spontaneous voltage oscillations in neuronal ensembles play a critical role in memory formation and storage. Although oscillatory activities arising endogenously within central pattern-generating (CPG) networks underlie many rhythmic motor behaviors, the contribution of such autonomous signals to motor learning and memory remains poorly understood. Previously, we found that the buccal CPG network driving food-seeking behavior in *Aplysia* contains a subset of electrically-coupled neurons that produces spontaneous, variable-amplitude voltage oscillations instigating infrequent and irregular cycles of patterned motor output. This pattern-initiating activity originates from organelle-derived, inositol triphosphate (IP3) receptor-dependent calcium oscillations in a pair of identified decision-making neurons (B63) within the CPG subset (Bédécarrats et al., 2021). Here, we show that the cycle frequency of this spontaneous pacemaker mechanism based on intracellular calcium store release is persistently increased by operant reward-learning, and in association with increased B63 excitability, constitutes a fundamental memory trace for accelerated and stereotyped rhythmic food-seeking movements.

**eLife Assessment:** Long-lasting plasticity in the dynamics of oscillatory intracellular calcium store release in key neurons of *Aplysia*’s food-seeking network has been identified as a novel sub-cellular substrate for operant-reward learning.

## Introduction

Oscillatory electrical activities occurring at frequencies below 1 Hz and arising in the absence of external stimuli are generated by a variety of central neuronal networks (Ahrens et al., 2013; Fenk et al., 2024; Gonzalo Cogno et al., 2024; Hanson, 2021; Yuste et al., 2024, 2005). A growing body of evidence from cortical neuronal assemblages suggests that such spontaneous, slow voltage oscillations may play a critical role in memory formation and the long-term storage of past behavioral experiences (Klinzing et al., 2019; Mölle and Born, 2011). In support of this notion, learning can persistently modify synchronized cortical oscillatory activities, and experimental manipulations of these rhythms modify long-term memory formation (Marshall et al., 2006; Miyamoto et al., 2017; Mölle et al., 2009, 2004; Ngo et al., 2013). A causal relationship between slow network oscillations and the formation and storage of memory engrams has also been found in sub-cortical and cortical regions associated with declarative memory, as well as in circuits involved in motor learning and non-declarative memory (Lemke et al., 2021; Nicolas et al., 2025; Sale and Kuzovina, 2022).

The membrane and synaptic properties underlying the autonomous genesis of slow network voltage oscillations have been extensively studied in motor CPG networks of vertebrates and invertebrates alike (Büschges and Ache, 2025; Calabrese, 1995; Mantziaris et al., 2020; Marder et al., 2005; Sakurai and Katz, 2015; Steuer and Guertin, 2019). These cellular properties can also be modified by learning paradigms and participate in memory storage (Baxter and Byrne, 2006; Benjamin et al., 2000; Hill et al., 2015; Lyons et al., 2005; Mueller et al., 2025; Namiki et al., 2024; Rivi et al., 2021). In addition to the contributions of synaptic and neuronal membrane properties, network voltage oscillations can arise from spontaneous organelle-driven fluxes in cytosolic calcium (Jackson and Thayer, 2006; Kikuta et al., 2019) that in turn may activate plasma membrane channels underlying impulse bursting (Bédécarrats et al., 2021). However, the involvement of such slow intracellular oscillatory processes in memory formation and storage, as well as the consequences for motor output production, remain largely unknown.

Here, we have addressed this issue in the marine mollusk *Aplysia*, in which various parameters of feeding behavior, including exploratory and consummatory biting movements of the tongue-like radula, can be modified by operant-reward conditioning (Brembs et al., 2002; Hawkins and Byrne, 2015; Lorenzetti et al., 2006; Nargeot et al., 2007; Susswein et al., 1986). Multiple forms of synaptic and membrane plasticity induced by such appetitive learning have been identified in the central neuronal circuitry that generates the buccal motor patterns (BMPs) that drive radula biting behavior (Baxter and Byrne, 2006; Brembs et al., 2002; Costa et al., 2020, 2022; Lorenzetti et al., 2008; Momohara et al., 2022; Mozzachiodi et al., 2008; Nargeot et al., 2009). In one type of operant learning, a contingent association for 30-40 min of a food reward with spontaneous biting movements transforms for several hours otherwise infrequent and sporadic exploratory biting into an accelerated and regularly recurrent expression that resembles other appetitive reward-induced patterns of compulsive behavior (Nargeot et al., 2007). Since the neural correlates of these learning-induced changes *in vivo* continue to be expressed by the buccal CPG circuitry in isolated CNS preparations, thereby allowing electrophysiological analysis at the cellular level, several neuronal and synaptic mechanisms contributing to this behavioral plasticity have already been elucidated (Nargeot et al., 2009; Sieling et al., 2014).

Recently, a subset of gap junction-coupled neurons belonging to the buccal CPG network was found to spontaneously produce synchronous membrane potential oscillations. This continuous low and variable amplitude rhythmic signal arises uniquely from a bilateral pair of decision making constituents of this subcircuit (the two B63 interneurons), originating from the dynamics of intracellular calcium release/re-uptake by internal organelles via inositol triphosphate (IP3) receptor activation. The resulting rhythmic cycles of plasma membrane depolarization/hyperpolarization can remain below the B63 neurons’ excitability threshold or, at irregular intervals, may trigger plateau potential-mediated bursts of action potentials that are necessary and sufficient to initiate individual BMPs and resultant radula bites (Bédécarrats et al., 2021; Hurwitz et al., 1997; Nargeot and Simmers, 2012). The present study shows that operant reward learning accelerates these organelle-derived, calcium-dependent voltage oscillations and that this novel form of plasticity, together with a decrease in the B63s’ excitability threshold, makes a critical contribution to the memory engram for a long-lasting transition to compulsive food-seeking action.

## Results

### Operant conditioning-induced regularization and acceleration of radula biting, and underlying BMP genesis *in vitro*

To investigate the effects of operant conditioning on the oscillatory mechanisms responsible for generating radula biting movements, two groups of animals (‘Contingent’ and ‘Non-contingent’) were trained using a previously described operant learning paradigm (Figure 1A; Nargeot et al., 2007). During a 30 min training period, all animals were exposed to a constant, non-ingested inciting stimulus applied to the lips to promote food-searching and radula biting behavior. In the Contingent group (n = 18), a food reward was delivered in association with each bite cycle, thereby establishing the action-reward contingency representative of operant conditioning. In the Non-contingent control group (n = 18), the same number of food rewards was delivered as for corresponding Contingent animals, but at fixed time intervals and independently of the animal’s ongoing biting behavior. Comparisons between these groups then allowed identification of the behavioral changes specifically induced by the action-reward association that characterizes operant conditioning. These changes were assessed during a 10 min test period, conducted 1 hr after training, during which time the animals were continuously exposed to the inciting stimulus alone. As expected from this learning paradigm, both the frequency and regularity of radula movement cycles were significantly higher in the Contingent group compared to the Non-contingent group (Figure 1B,C, left *vs.* right; D1,D2; Rate: W = 291.5, p <0.001; Coefficient of variation: W = 51.5, p < 0.001). These findings are therefore consistent with previous reports that operant conditioning leads to an acceleration and regularization of *Aplysia’s* food-seeking actions, and these behavioral changes persist for several hours following contingent training (Nargeot et al., 2007, 2009).

**Figure 1.**
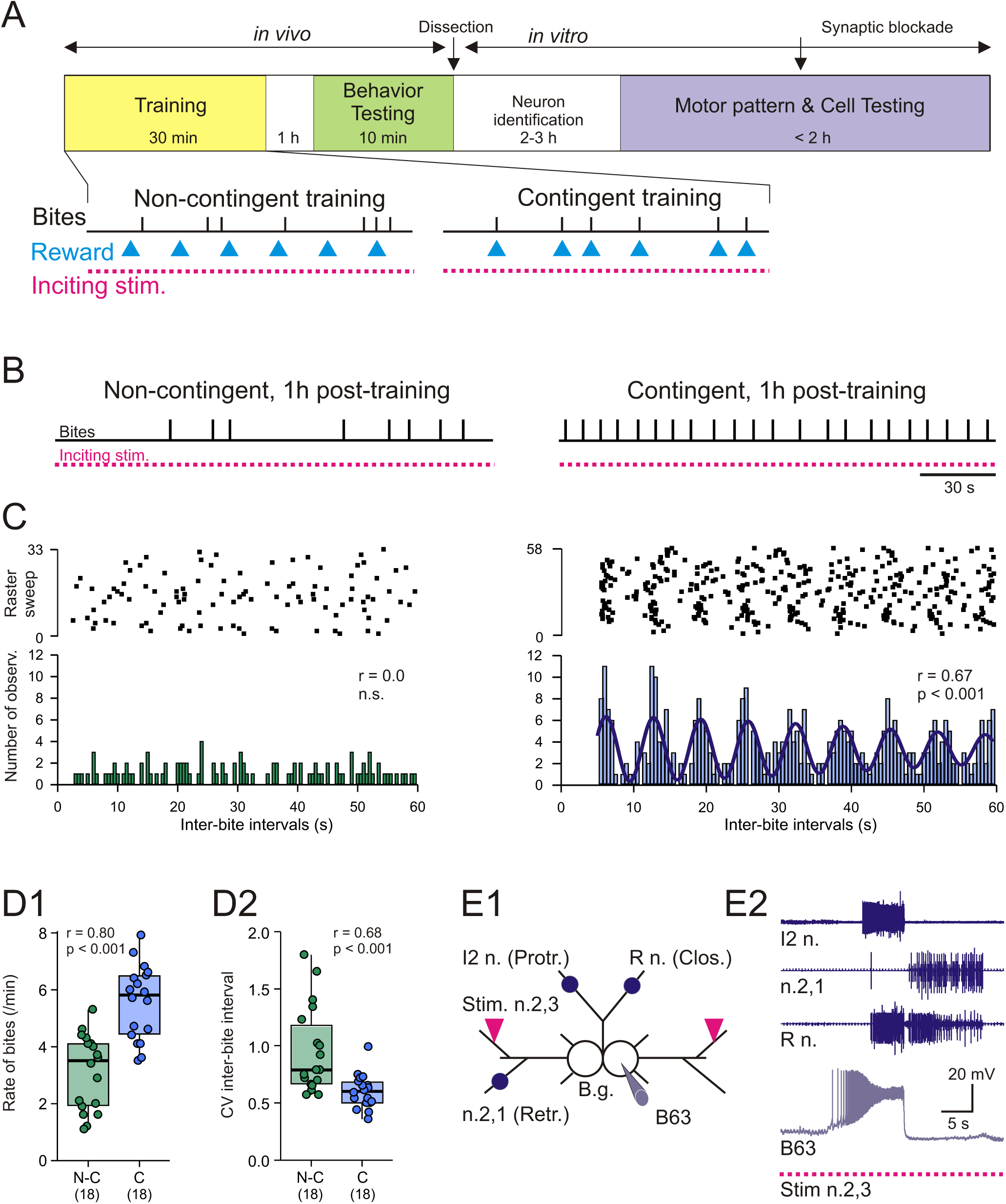
Behavioral plasticity induced by operant conditioning and the *in vitro* buccal ganglia preparation. (**A**) Experimental paradigm: The initial *in vivo* protocol consisted of training (yellow box) and subsequent testing (green box) phases. During the training period (bottom traces), animals in a ‘Contingent’ group (at right) were continuously stimulated with an inciting food stimulus (dashed red lines) and received a food reward (blue arrowheads, right) in association with each spontaneous radula bite cycle (vertical bars indicate the onset of each movement cycle), thereby reproducing the action-reward contingency characteristic of operant conditioning. Animals in a different control ‘Non-contingent’ group (at left) received the same inciting stimulus and the same number of food rewards, but delivered at fixed time intervals, independently of the animal’s biting behavior (blue arrowheads, left). Neuronal changes induced by these *in vivo* training paradigms were subsequently assessed in isolated buccal ganglia *in vitro* (see E1) during a ‘Motor & Cell testing’ period (purple). (**B**) Representative excerpts of biting behavior recorded 1 hr after Non-contingent (left) and Contingent (right) training in the presence of an inciting food stimulus. **C)** Raster plots (top) and corresponding autocorrelation histograms (bottom), of the interval between successive bites in different Non-contingently- and Contingently-trained *Aplysia* during the 10 min testing period. The Non-contingent animal (left) expressed infrequent and irregular bite cycles, whereas the Contingent animal (right) displayed regularly rhythmic biting, as indicated by a significant Gabor function fit to the autocorrelation histogram (bold sinusoidal line, *p* < 0.001). **(D)** Group quantification of biting behavior during the post-training test period: (D1) bite cycle rate and (D2) coefficient of variation of inter-bite intervals. Operant conditioning significantly increased both the frequency and regularity of radula movement cycles. Numbers in brackets in this and the following figures indicate numbers of animals per group. **(E)** Left: Schematic of the isolated buccal ganglia preparation (E1) showing positions of recording electrodes (blue dots) placed on selected motor nerves, and stimulating electrodes (red triangles) placed on the bilateral sensory nerves n.2,3 to promote buccal CPG activation. (E2) Extracellular recordings of an individual buccal motor pattern (BMP; top three traces) that *in vivo* would produce a single radula bite cycle, and simultaneous intracellular recording of the associated action potential burst and underlying plateau potential in the B63 neuron of the right buccal ganglion (bottom trace). The fictive bite consists of protraction, retraction and closure burst phases (recorded from motor nerves l2 n., n.2,1 and R n., respectively), which are triggered in the buccal CPG network by plateau-driven impulse bursts in the B63 neurons during inciting nerve 2.3 stimulation (Nargeot et al., 2007).

The neuronal correlates of this learning-induced plasticity expressed by these animals were next analyzed in their still active buccal ganglia after isolation *in vitro* and maintenance under artificial sea water (ASW; Figure 1E1). Simultaneous recordings of the motor output responsible for biting behavior, which consists of repeating cycles of a two-phase-buccal motor pattern (BMP; Figure 1E2) that drives radula protraction, closure and retraction, and associated variations in the membrane potential of one or both bilateral B63 neurons were then performed during tonic stimulation of input nerve 2,3 (n.2,3; 0.3 ms pulse, 8 V, 2 Hz). This latter sustained stimulation, which was applied over a 20 min test period, served as an *in vitro* analog of the inciting sensory stimulation previously applied *in vivo* (Nargeot et al., 2007). Recordings were analyzed starting 10 min after the onset of afferent nerve stimulation in two subgroups of isolated preparations (n = 9 per subgroup) randomly assigned from the two main groups according to their previous Contingent *vs.* Non-contingent training.

Consistent with radula biting movements after operant conditioning in the intact animal, both the rate and regularity of BMP production remained significantly increased in the first Contingent subgroup compared with the Non-contingent subgroup of ganglia (Figures 2A1,B1; 3A,B Rate: H = 26.484, p < 0.001; q = 2.176, p = 0.044; Coefficient of variation computed in preparations that exhibited at least three BMPs: H = 16.42, p < 0.001; q = 2.527, p < 0.023). These results therefore corroborate the earlier finding that the buccal CPG network in isolated buccal ganglia under continuous inciting stimulation retains, for several hours, fundamental correlates of the behavioral plasticity induced by prior operant conditioning in the intact animal (Brembs et al., 2002; Nargeot et al., 2007, 2009).

**Figure 2.**
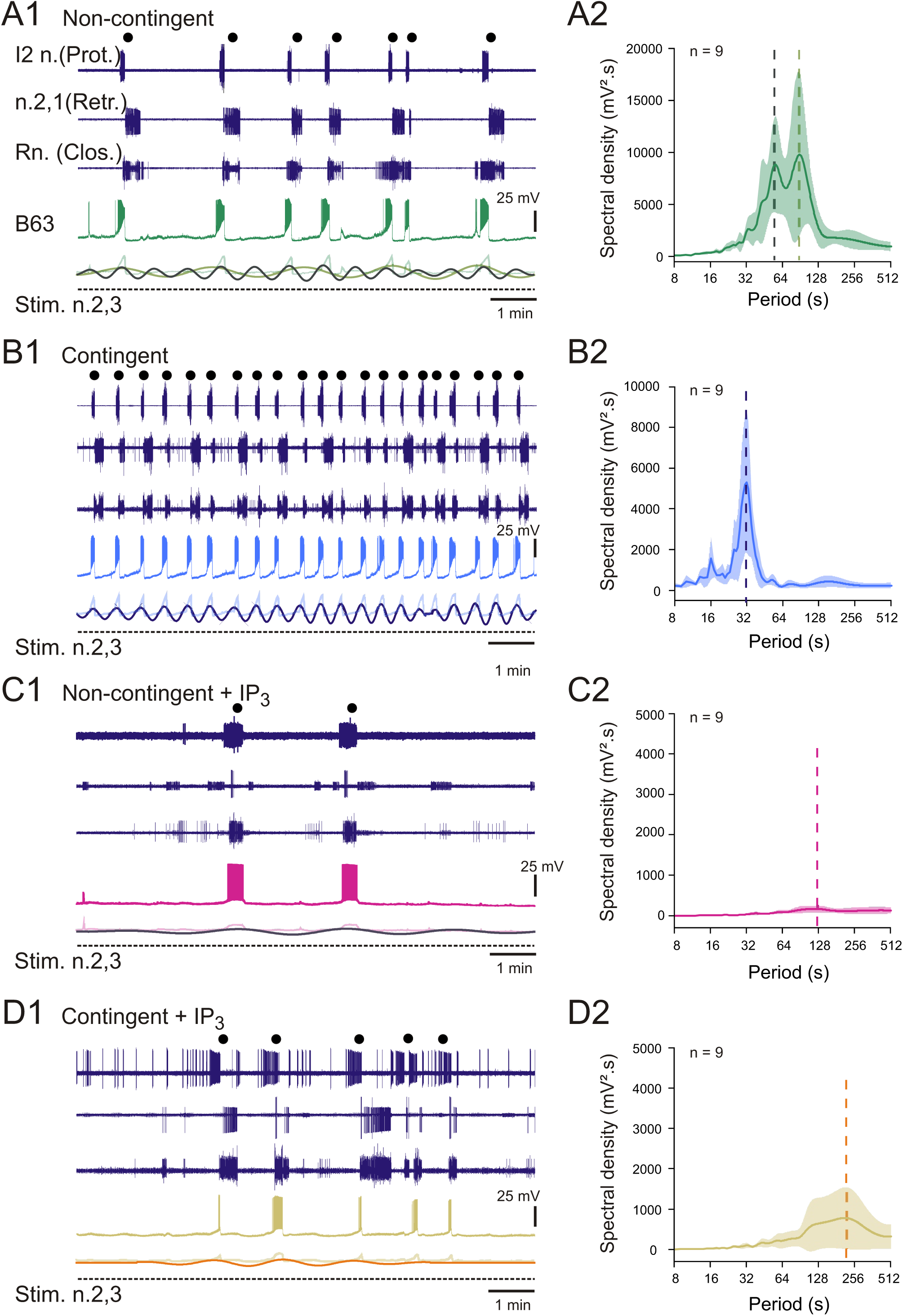
Learning-induced changes in BMP generation and associated IP3-sensitive oscillatory activity of B63 neurons in isolated buccal ganglia under input nerve stimulation. (**A**) Data from *in vitro* preparations isolated from Non-contingent animals (n = 9 in a randomly assigned subgroup of N-C animals assessed in Figure 1D): A1, representative 10 min recording of BMPs (black dots) recorded in buccal motor nerves (upper three traces) and underlying membrane potential variations in a B63 neuron during tonic stimulation of nerve 2,3 (n.2,3; 8 V, 0.3 ms-pulse, 2 Hz, dashed line). During this inciting stimulation, the B63 cell generated infrequent and irregular bursts of action potentials and resultant BMPs associated with periodic variations in the neuron’s membrane potential. Lower sinusoidal waveforms: wavelet-based reconstructions of the two dominant rhythms decomposed from B63’s membrane potential fluctuations associated with the cell’s bursting and BMP production, superimposed on the smoothed raw recording trace (faded trace; see Figure 2-figure supplement 1 for details). A2, averaged density plot (+/-CI_95%_) from spectral analyses of B63 neuron recordings in all 9 Non-contingent preparations, showing the two distinct periodicities (dashed vertical lines) in the cell’s voltage fluctuations. Note that the dashed lines in the averaged FFT indicate the mean power density peaks of the two dominant rhythms (i.e., largest averaged amplitudes), rather than their median cycle periods. **(B)** Equivalent presentation as in A, but for preparations from Contingently trained animals (n = 9 in a randomly assigned subgroup of Contingent (C) animals assessed in Figure 1D) during the same tonic n.2,3 stimulation. B1, such a preparation expressed more frequent and regular BMPs in association with an accelerated suprathreshold oscillation of B63’s membrane potential (bottom colored sinusoid). B2, averaged spectral density plot (+/-CI_95%_) from all 9 Contingent preparations indicating that B63’s voltage fluctuations were now composed of a solitary dominant period (dashed vertical line) correlated 1:1 with each B63 burst and resultant BMP (see B1). **(C)** Non-contingent preparations (n = 9 from the remaining subgroup of N-C animals; see Figure 1D) with the bilateral B63 neurons iontophoretically injected (for 1 hr) with inositol triphosphate (IP3, 10 mM) and subjected to tonic n.2,3 stimulation. C1, In a representative example, BMPs were now sporadically expressed in association with diminished, low frequency fluctuations in B63’s membrane potential (bottom colored sinusoid); C2, averaged spectral density plot (+/-CI_95%_) from this Non-contingent subgroup. **(D)** Contingent preparations (n = 9 from the remaining C animals; see Figure 1D) with IP3-injected B63 neurons (10 mM for 1 hr). D1, Tonic input nerve stimulation no longer elicited accelerated and regularized buccal motor output (cf. B1), but as for IP3-injected N-C preparations, led to erratic BMPs associated with suppressed, low frequency fluctuations in B63’s membrane potential (lower colored sinusoid). D2, averaged spectral density plot (+/-CI_95%_) from this Contingent subgroup.

### The learning-induced plasticity is associated with changes to an IP3-sensitive voltage oscillation in the B63 neurons

The production of individual BMPs driving each radula bite cycle arises from spontaneous rhythmic variations in membrane potential of the two electrically-coupled B63 neurons, which that in turn instigate BMP genesis by the wider buccal CPG network (Bédécarrats et al., 2021). These cyclic, variable-amplitude B63 depolarizations can remain subthreshold for impulse generation or, at irregular intervals, may trigger a large-amplitude plateau potential depolarization accompanied by an intense burst of action potentials and the expression of a BMP (see Figure 2A1). In Non-contingent preparations, Fourier (spectral) analyses (Figure 2A2) combined with wavelet reconstructions show that these voltage fluctuations decompose into two distinct dominant periodicities (Figure 2A1, lower traces; Figure 2-figure supplement 1A): a slower rhythmic signal (median period 86.9 s) that is correlated with the plateau potential-driven impulse bursts and resulting BMPs, and embedded in a faster oscillatory waveform (median period 52.1 s) whose depolarized peaks are only intermittently associated with B63 bursts and BMP production. The lower frequency signal is associated with the expression of B63s’ voltage-dependent membrane properties (plateauing and bursting) and the occurrence of 0, predominantly 1, or occasionally 2 BMPs per oscillation cycle depending on buccal network excitability. On the other hand, the faster voltage waveform presumably arises from an oscillatory mechanism that involves an intracellular calcium dynamic requiring IP3-receptor activation and whose frequency is independent of B63’s membrane potential (Bédécarrats et al., 2021; also see below).

Operant conditioning modified the frequency of these B63 oscillations under the tonic inciting stimulation, with the cycle periods of both the faster and slower voltage waveforms being reduced in the Contingent group (n = 9) compared with the Non-contingent (n = 9) preparations (Figure 2B1,B2). This dual frequency increase was such that the two voltage waveforms were now coordinated in synchrony, as indicated by the single dominant peak (median period 29.8 s) in the averaged periodogram of Figure 2B2 (see also Figure 2-figure supplement 1B), with a B63 burst and associated BMP occurring virtually 1:1 with the depolarized crest of each accelerated fast rhythm cycle (Figure 2B1). In contrast to this increase in frequency of the faster voltage oscillation (Figure 3C; H = 26.754, p < 0.001; Contingent vs. Non-contingent: q = 2.105, p = 0.042), learning did not result in a significant modification to the oscillation’s magnitude (Figure 3D; H = 24.892, p < 0.001; Contingent vs. Non-contingent: q = 0.671, p = 0.531).

**Figure 3.**
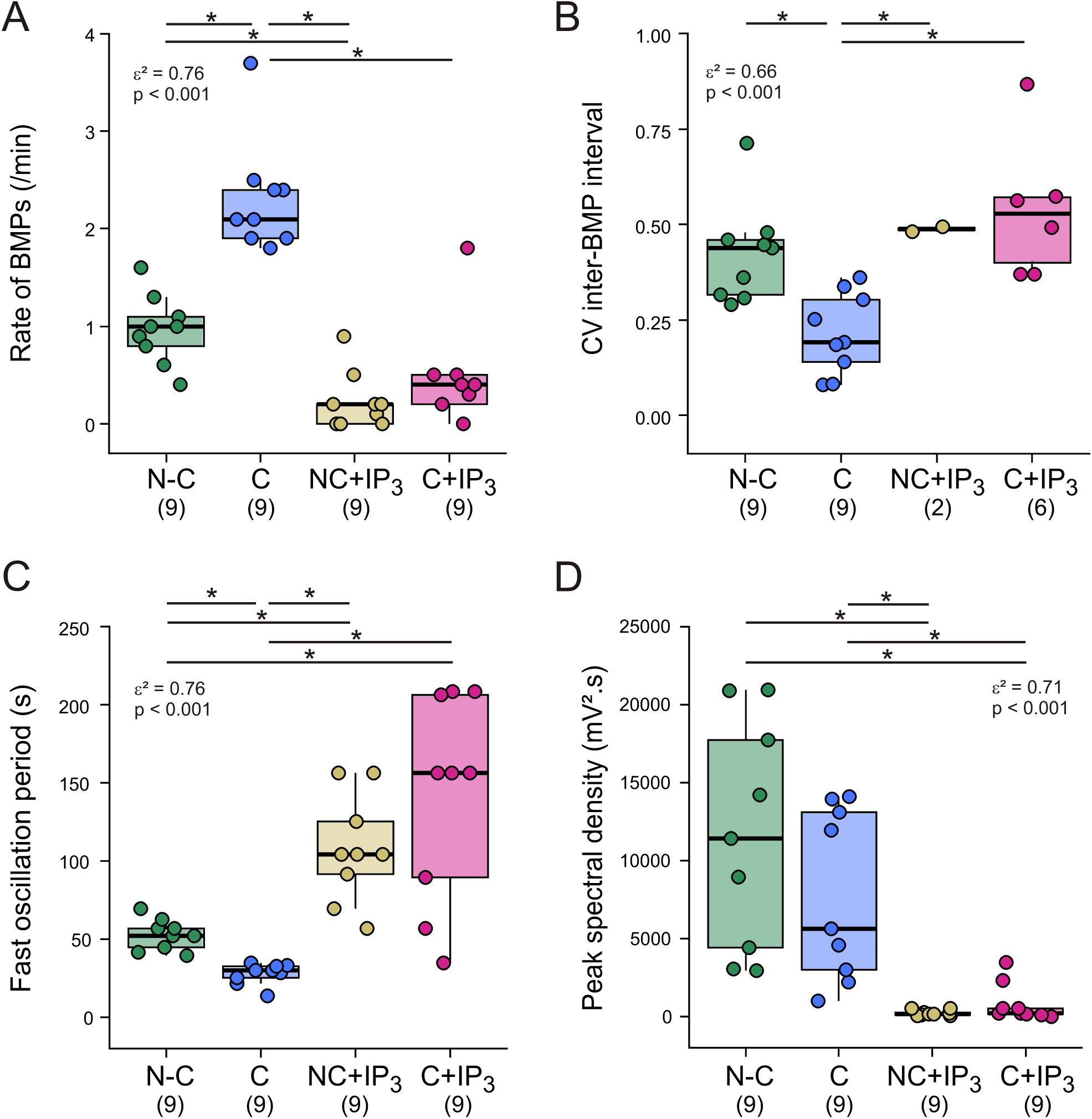
Quantitative analysis of the plasticity in BMP genesis and B63’s voltage oscillations expressed during input nerve stimulation. (**A,B**) Group comparisons of BMP production in a 10 min-test period between the four experimental conditions as illustrated in Figure 2. After prior operant conditioning *in vivo*, the frequency (A) and regularity (B) of motor pattern cycles *in vitro* increased significantly in the Contingent (C) group compared with the Non-contingent (N-C) group, but both parameters were decreased in the two groups containing IP3-injected B63 neurons (NC+IP3, C+IP3). Note that the coefficient of variation of inter-BMP intervals (B) was reported only when at least three BMPs occurred during the test period, consequently excluding 7/9 Non-contingent and 3/9 Contingent preparations, both with IP3-filled B63 neurons, from this analysis. (**C,D**) Group comparisons of the cycle period (C) and amplitude (D) of the faster dominant voltage oscillation of B63 neurons in the same preparations as in A,B (including those in Figure 2). The oscillation frequency increased (i.e. period decreased, C) with no change in amplitude (D) in the Contingent group compared with the Non-contingent group, but both oscillation parameters were strongly reduced in the two groups with IP3-filled B63 neurons.

To assess the involvement of an IP3-sensitive pacemaker mechanism in the learning-induced plasticity of BMP genesis, we used injections of exogenous IP3 to experimentally modify cytosolic levels of this second messenger in the B63 neurons. Accordingly, for the second randomly assigned Non-contingent subgroup (n = 9; Non-contingent+IP3) and the second Contingent subgroup (n = 9; Contingent+IP3) that had previously undergone the same *in vivo* training protocols as described above, both bilateral B63 neurons were iontophoretically injected with an IP3-potassium solution (10 mM; pulses: -1 nA, 0.5 s, 1 Hz) for 1 hr. Such injections, and subsequent recording in the presence of the IP3-containing electrode were used to maintain intracellular IP3 levels at an elevated level, a condition expected to alter IP3 receptor gating and consequently suppress the B63s’ organelle-derived calcium oscillations (Hajnóczky and Thomas, 1997; Wakui et al., 1989, also see Discussion).

In both the Non-contingent and Contingent groups with IP3-loaded B63 neurons, the rate and regularity of buccal motor cycles were reduced compared with the equivalent groups without IP3 loading (Figures 2C,D; 3A,B; Rate: Contingent+IP3 vs. Contingent: q = 3.960, p < 0.001; Non-contingent+IP3 vs. Non-contingent: q = 2.546, p = 0.022; Coefficient of variation evaluated when at least 3 BMPs were expressed, Contingent+IP3 vs. Contingent: q = 3.652, p = 0.002; Non-contingent+IP3 vs. Non-contingent: q = 1.022, p = 0.368). The lack of an evident significant difference in BMP regularity between these latter two groups was most probably due to the small number of Non-contingent preparations (n = 2 from 9) that expressed >3 BMP cycles after injection of IP3. Furthermore, in correlation with motor output changes and in addition to the decreased frequency of their voltage oscillation, the amplitudes of oscillation in IP3-loaded B63 neurons in both Non-contingent and Contingent preparations were drastically reduced (Figure 2C,D, Figure 2-figure supplement 1C,D).

The finding that intracellular IP3 injection reduced the oscillation frequency of B63 in both Contingent and Non-contingent ganglia compared to their non-injected counterparts (Figure 3C; Contingent+IP3 vs. Contingent: q = 4.512, p < 0.001; Non-contingent+IP3 vs. Non-contingent: q = 2.161, p = 0.042), therefore indicated that the modulator’s effect on B63’s oscillatory properties was not specific to a paradigm of prior *in vivo* training. Indeed, that no significant difference in the frequency of the residual oscillations remained between the Non-contingent and Contingent preparations indicated that the elevated presence of IP3 had additionally obscured the memory trace of operant learning (Figure 3C; Contingent+IP3 vs. Non-contingent+IP3: q = 0.246, p = 0.805; see Discussion). Finally, whereas the amplitudes of voltage oscillations in non-injected B63 were not modified by learning (see above), they were strongly and similarly decreased following IP3 injection in Contingent and Non-Contingent preparations, and therefore independently of learning (Figure 3D; H = 24.892, p < 0.001; Contingent+IP3 vs. Contingent: q = 2.819, p = 0.007; Non-contingent+IP3 vs. Non-contingent: q = 4.116, p < 0.001; Contingent+IP3 vs. Non-contingent+IP3: q = 0.626, p = 0.531).

Overall, these results indicate that the frequent and regular, compulsive-like buccal motor pattern expression occurring after learning, originates from an accelerated, low-amplitude voltage oscillation in the decisional B63 neurons. This oscillatory activity is sensitive to the intracellular IP3 content of the cells, independently of the prior training protocol, thus further indicating the second messenger’s crucial importance to the basic oscillatory mechanism. Moreover, the finding that elevated IP3 levels extinguish the learning-induced plasticity by reducing the oscillation frequency of Contingent IP3-injected B63 neurons to levels similar to those observed in Non-contingent IP3-loaded cells suggests an important role for IP3 receptor dynamics in the actual learning process.

### The changes in B63’s voltage oscillation arise independently of CPG network inputs

In principle, the learning induced plasticity in B63’s membrane potential oscillation and resultant BMP production could derive from instructive influences that are intrinsic to buccal CPG circuitry and/or are conferred by long-lasting modifications in synaptic pathways that are upstream to the network, including those conveying sensory information from the periphery. To assess this latter possibility, therefore, BMP genesis and the accompanying variations in B63’s membrane potential were subsequently recorded in isolated buccal ganglia in the absence of any input nerve stimulation. Recordings of this strictly spontaneous CPG activity were performed during a 10 min test period beginning after an equivalent post-training delay in the subgroups (n = 9 per subgroup) of Non-contingent, Contingent, and Contingent+IP3-injected B63 preparations (mean delay: Non-contingent, 5.3 hr; Contingent 5.4 hr; Contingent+IP3, 5.2 hr; 3-group comparison: H = 1.062, p = 0.598). Since the Non-contingent and Contingent preparations with IP3-injected B63 neurons showed no significant differences in terms of either BMP production or B63 oscillatory activity, only the Contingent+IP3 group was further investigated in detail.

In the absence of input nerve stimulation, and regardless of the experimental group, BMPs continued to be generated, albeit at expected lower overall frequencies than those elicited during inciting input nerve stimulation (Figure 4A1, B1). However, both the rate and regularity of these spontaneous BMPs remained significantly higher in ganglia from contingently trained than from non-contingently trained animals (Figures 4A,B; 5A,B: Rate: W = 66, p = 0.026; CV computed from preparations exhibiting at least three spontaneous BMP cycles: W = 5.5, p = 0.020), whereas no BMPs were generated in the Contingent+IP3 preparations (Figure 4C). Thus, these new findings indicate that the memory trace produced by operant conditioning is not stored in sensory input or other extrinsic pathways that affect BMP genesis when activated. Instead, the memory is inscribed in the intrinsic functional dynamics of the central circuitry that autonomously generates radula biting movements.

**Figure 4.**
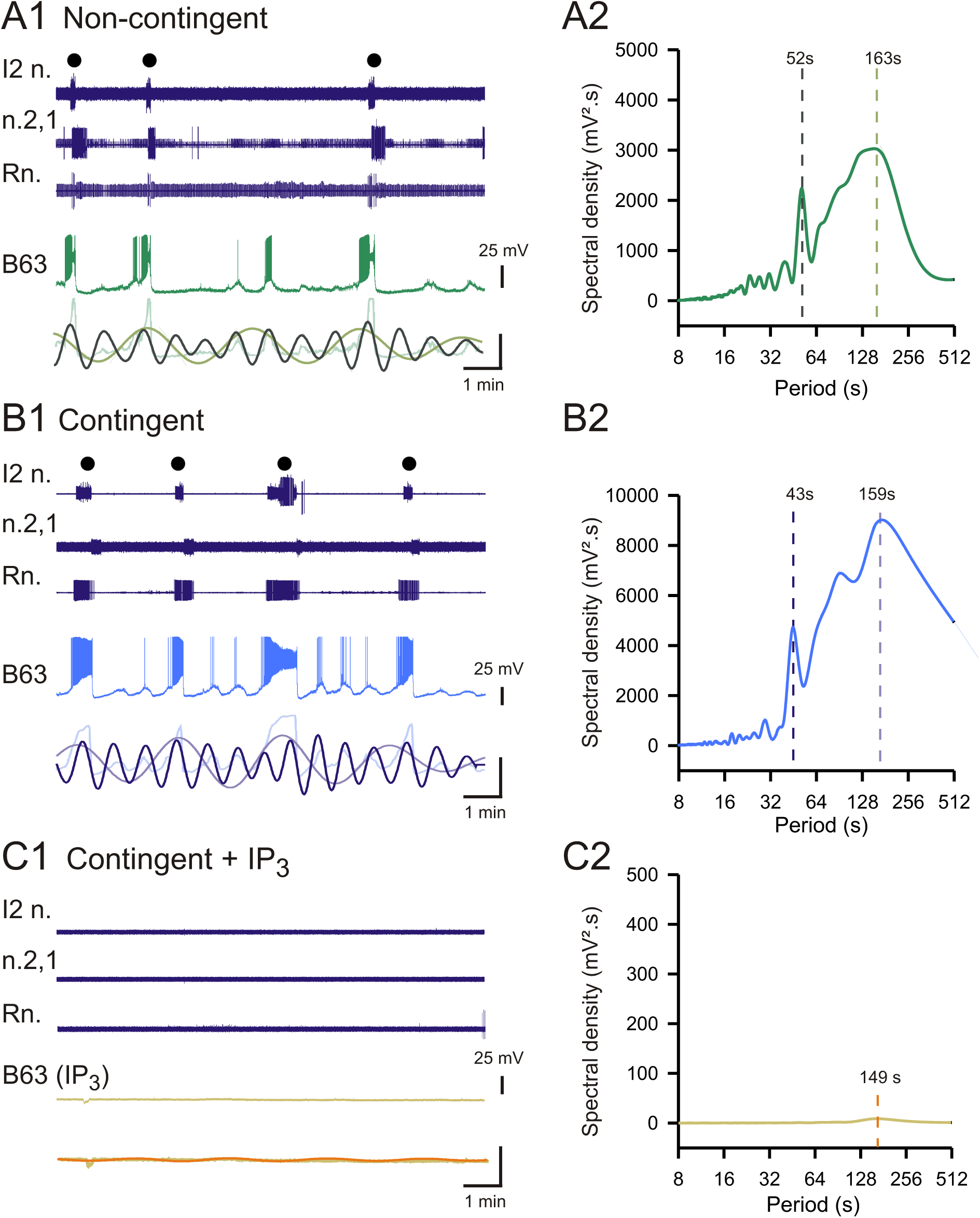
Learning-induced changes in spontaneous BMP generation and associated B63 voltage oscillations in the absence of input nerve stimulation. (**A1-C1)** 10-min recording excerpts of spontaneous B63 activity and associated BMP production (top three traces, black dots) in different isolated ganglia: Non-contingent (A1), Contingent (B1) and a Contingent preparation with IP3-loaded B63 neurons (C1). (A2-C2) Corresponding spectral density plots of the B63 cells recorded in A1-C1. The spontaneous voltage fluctuations of non-injected B63 neurons decomposed into two dominant periods (A2,B2, dashed lines) whose wavelet-based reconstructions, when superimposed on the smoothed B63 voltage recordings (bottom traces in A1,B1), correlated with the expression of large amplitude plateaus and resultant BMPs. These spontaneous voltage oscillations were slower in the Non-contingent preparation (A) than in the Contingent preparation (B), whereas the single oscillation that now remained subthreshold for plateauing and BMP production was almost suppressed in the Contingent preparation with IP3-filled B63 neurons (C).

As described above, the spontaneous production of BMPs is correlated with a slow rhythm associated with impulse bursts and plateau potentials in the B63 neurons which is coordinated to a faster, low-amplitude rhythmic signal that is timed with all membrane voltage changes (also see Bédécarrats et al., 2021). Spectral analysis and associated reconstructions of these waveforms indicated that their frequency was different between the experimental groups. In correspondence with the increased rate of BMP production, the slower B63 rhythm was expressed at a higher frequency in the Contingent preparations than in the Non-contingent preparations (Figure 4A1,A2; B1,B2; W = 9, p = 0.039), and was absent in the Contingent+IP3 loaded preparations (Figure 4 C1,C2; Figure 4-figure-supplement 1A-C). Similarly, operant conditioning modified B63’s faster oscillatory rhythm, with its median cycle period being significantly reduced in the Contingent group compared with the Non-contingent group, whereas it increased in the Contingent+IP3 group (Figure 5C; H = 15.374, p < 0.001; Contingent vs. Non-continent: q = 1.961, p = 0.05; vs. Continent+IP3: q = 3.921, p < 0.001; Non-continent vs. Continent+IP3: q = 1.961, p = 0.05). However, this learning-induced acceleration was not associated with a significant difference in oscillation amplitude in Contingent and Non– contingent preparations, with both waveforms remaining considerably larger than the residual signal expressed in Contingent preparations with IP3-injected neurons (Figure 5D; faster rhythm, H = 17.802, p < 0.001, Contingent vs. Non-contingent, q = 1.307, p = 0.191, vs. Contingent+IP3, q = 4.128, p < 0.001; Non-contingent vs. Contingent+IP3, q = 2.821, p < 0.007; slower rhythm: W = 29, p = 0.864).

**Figure 5.**
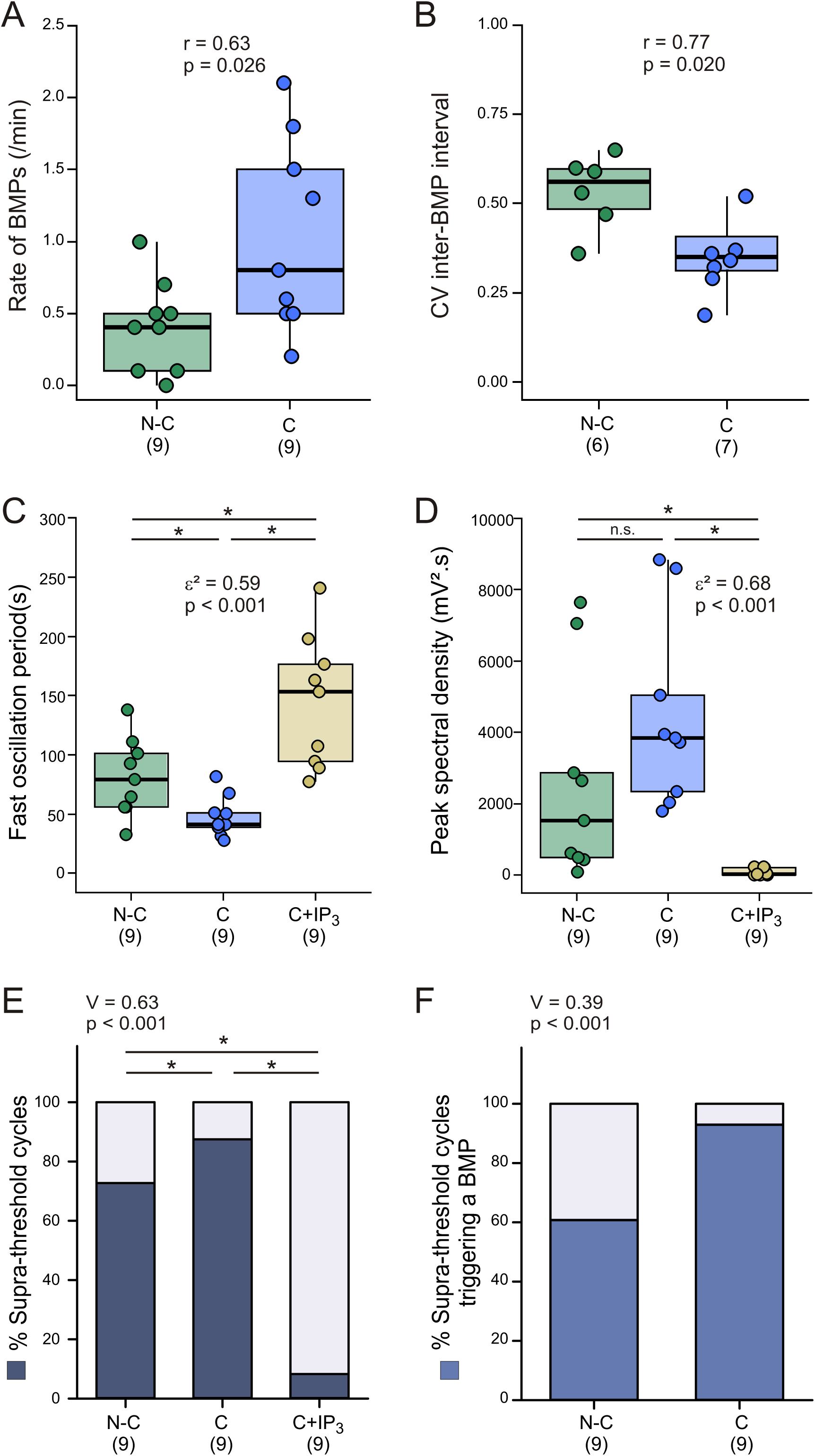
Quantification of the learning-induced plasticity in spontaneous BMP generation and in B63’s underlying voltage oscillations and excitability. (**A,B**) Between group comparisons of the same isolated preparations as illustrated in Figure 4 (n = 9 per group) showing that without tonic input nerve stimulation, both the frequency (A) and regularity (B) of spontaneously produced BMPs increased significantly in ganglia from contingently trained animals compared with Non-contingent ganglia. No BMPs were expressed in Contingent preparations with IP3 loaded B63 neurons. Note that CV values could be computed from 6/9 Non-contingent and 7/9 Contingent preparations that expressed at least 3 BMPs. (**C,D**) Group comparisons of the cycle period (C) and amplitude (D) of the faster dominant oscillation in B63 neurons, expressed in the same isolated preparations as in A,B. Prior i*n vivo* contingent training (middle boxes) significantly increased the frequency (i.e., period decreased, C), with no significant change in amplitude (D) of this spontaneous, low amplitude oscillation compared with Non-contingent controls (left boxes). Both the waveform’s frequency and amplitude were markedly reduced in Contingent preparations with IP3-injected B63 neurons (right boxes). (**E,F**) Between group comparisons of the proportion of faster B63 oscillation cycles that triggered action potentials (E, dark blue shaded areas), and of these supra-threshold cycles, those that initiated a plateau potential and a resultant BMP (F). Both proportions were significantly higher in Contingent ganglia than in either the Non-contingent or Contingent+IP3 loaded groups. Note that in this latter group, very few oscillation cycles triggering action potentials were recorded (E, right box), and none of these led to BMP generation, so could not be quantified in F.

Despite this lack of learning-related changes in oscillation magnitudes, the proportion of faster cycles that elicited a burst of action potentials (defined as spike trains > 1 Hz lasting > 2 s) was significantly higher in the Contingent group than in the Non-contingent group (Figure 4A,B). This proportion was significantly reduced in the Contingent+IP3 group compared to the two other groups, with loaded B63 cells only rarely producing impulse firing (Figures 4C, 5E: Contingent vs. Non contingent, p = 0.013; Contingent+IP3 vs. Contingent, p < 0.001; vs. Non-contingent, p < 0.001). Moreover, among the supra-threshold B63 cycles expressed in Contingent and Non-contingent ganglia, the proportion of these cycles that triggered a plateau-like depolarization and resultant BMP was significantly higher in the Contingent group than in the Non-contingent group (p < 0.001), whereas plateauing activity was never observed in Contingent+IP3 preparations (Figures 4, 5F).

Overall, these results indicate that the learning-induced increase in the frequency and regularity of BMP production, in the absence of extrinsic stimulation, arise from an acceleration of spontaneous membrane potential oscillations in the B63 neurons, and by an increased likelihood that individual oscillation cycles trigger an impulse burst that drives a BMP cycle. This plasticity is drastically altered by the continuous presence of exogenous IP3, further indicating that it at least partly derives from learning-evoked changes to the organelle-related, calcium-dependent oscillatory mechanism within the B63 neurons themselves.

### The learning-induced acceleration is conferred by alterations to B63’s endogenous pacemaker mechanism

In addition to learning-related changes to an intrinsic, organelle-dependent pacemaker mechanism, the acceleration of the faster, low amplitude oscillation in the B63 neurons could also arise from a pre-synaptic source within the central network and/or from modifications to voltage-dependent properties of B63‘s plasma membrane channels. To experimentally dissociate these possibilities, buccal ganglia were bathed in a modified saline solution containing a low calcium concentration (3 mM) and the addition of cobalt (10 mM; ‘Low Ca^2+^ + Co^2+^ solution’), which was previously found to block chemical synapses (Bédécarrats et al., 2021).

Under these conditions, after an initial prolonged depolarization, recorded B63 cells then repolarized to initial membrane potential levels and continued spontaneously to express a low-amplitude oscillation with brief impulse firing, but now only occasionally accompanied by plateau depolarizations and associated prolonged burst discharge (Figure 6A). The reliance of this persistent oscillatory capability under functional synaptic isolation on an organelle-derived, IP3 receptor-sensitive mechanism was further confirmed by transiently injecting IP3 into the B63 neurons of a further control group of buccal ganglia preparations. Brief air-pressure pulses (5-10 PSI) of increasing duration (250-950 ms) delivering IP3 (10 mM) into a recorded B63 evoked individual depolarizing responses whose amplitude and duration progressively increased with the injection duration until impulse discharge occurred (Figure 6-figure-supplement 1). The effect of such experimentally-evoked depolarizations by injected IP3 was to cause a resetting of the phase of the ongoing B63 rhythm (Figure 6B). However, with longer pulse durations (> 750 ms), the pulsed IP3-evoked depolarization was followed by a slow return to the cell’s resting membrane potential, resulting in the next spontaneous voltage oscillation being transiently delayed (Figure 6C). Importantly, comparable injections of control solutions of potassium-acetate with Fast-Green (KAc, 10 mM; FG, 2 mM) or Fast-Green alone (10 mM) failed to exert any similar effects, thereby ruling out non-specific chemical or mechanical contributions to the IP3-evoked responses (Figure 6D; Figure 6-figure-supplement 1A,B). In contrast, when functionally isolated B63 neurons were subjected to intracellular injections of depolarizing current pulses of varying duration and amplitude (5-20 s, 1-5 nA), in a manner equivalent to the pulsatile IP3 injections, such imposed brief currents could neither reset nor suppress the ongoing voltage oscillation, consistent with a lack of a participation of voltage-dependent membrane channels in actual oscillation genesis (Figure 6E). Finally, it is noteworthy that whereas a 1 hr iontophoretic injection of IP3 caused a sustained suppression of B63’s rhythmic voltage signal as reported above, equivalent continuous injections of control solutions had no effect on the cell’s oscillation (Figure 6-figure supplement 2). Together, these results provide further evidence confirming that under conditions of chemical synaptic blockade, the persistent low amplitude oscillation occurring in B63 arises solely from IP3-sensitive, organelle-derived intracellular calcium dynamics without a significant contribution of voltage-dependent plasma membrane channels.

**Figure 6.**
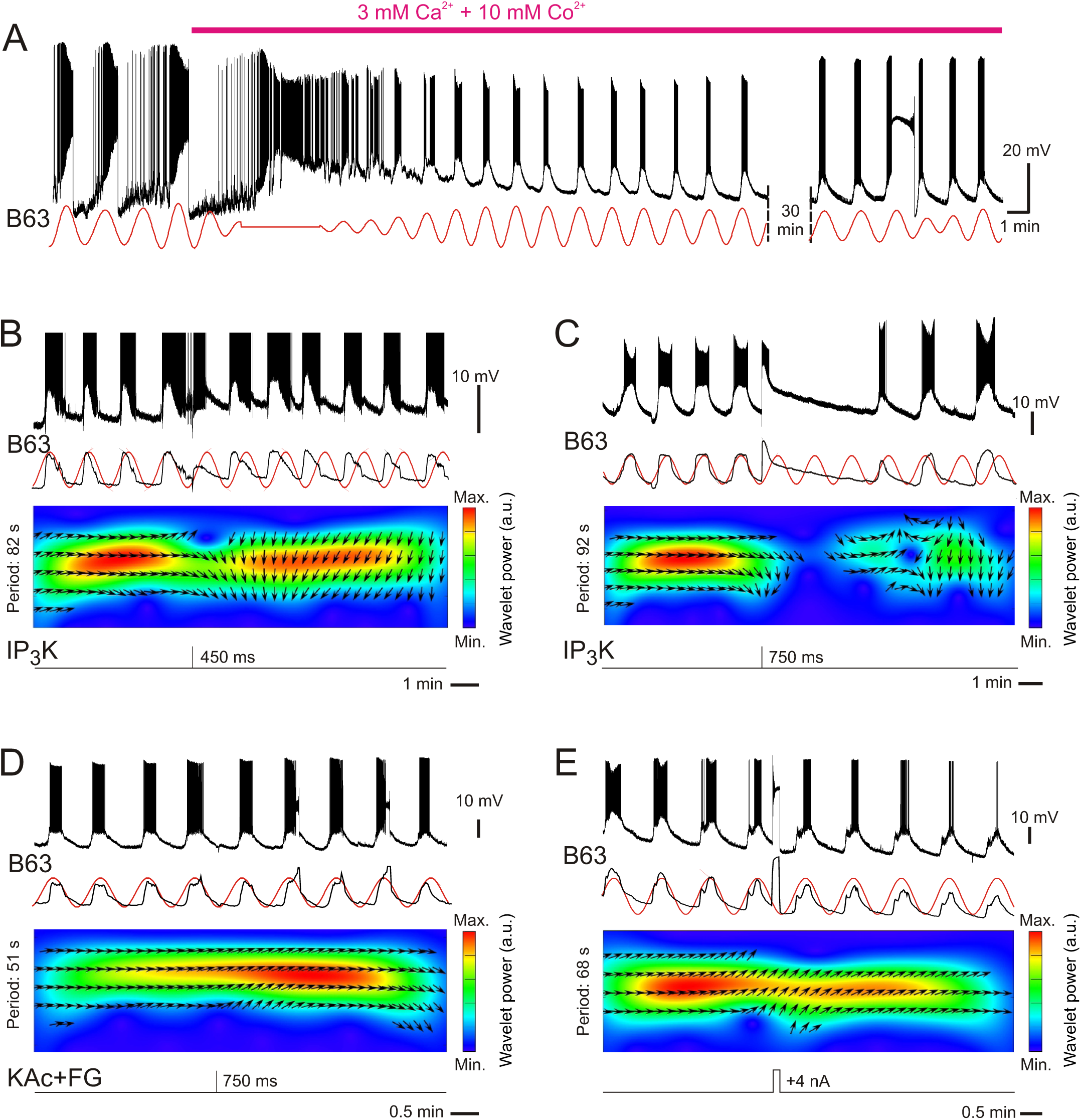
Transient effects of pressure-injected IP3 into B63 neurons under chemical synapse blockade. (**A**) Intracellular recording of a B63 neuron’s voltage fluctuations and corresponding reconstructed waveform from the peak spectral density (red sinusoid, period: 94 s) before and during bath-applied ‘low Ca^2+^ + Co^2+^‘ solution to block chemical synaptic transmission in the buccal CPG network. Under this condition, low-amplitude voltage oscillations continued to occur spontaneously and reliably trigger a burst of full-size action potentials in each cycle, occasionally eliciting a plateau potential characterized by a sustained depolarization underlying high- frequency, low-amplitude action potentials, presumably attenuated due to a negative interplay between the high firing rate of impulses and their relative refractory periods (see right). **(B,C)** Dose-dependent effects of intracellular IP3 (10 mM) on B63’s low-amplitude voltage oscillation in a different preparation under chemical synapse blockade as in A. A brief IP3K injection (pressure pulse: 450 ms, 5 PSI) transiently depolarized the cell, elicited a premature oscillation and impulse burst cycle and caused a phase advancement of the ongoing voltage oscillation (B; action potentials are partially truncated). Similarly, a longer pulse (750 ms, 5 PSI produced an initial depolarization and burst discharge, but with a subsequent slow repolarization and a delay of several minutes before resumption of the B63’s intrinsic oscillation (C). The phase shift was quantified by cross-wavelet analysis between B63’s membrane potential fluctuations (middle black traces: smoothed raw recordings) and a reference sinusoid (red) computed from the pre-pulse oscillation’s peak spectral density. The cross-wavelet power spectra (colored panels) show the time-resolved covariation of oscillation amplitude at the pre-pulse cycle frequency. Arrows indicate the instantaneous phase relationship (orientation towards right: in phase; towards left: out of phase; upwards: phase delay; downwards: phase advance). **(D,E)** Control experiments: Lack of responsiveness to an intracellular injection of a control solution potassium acetate (KAc, 10 mM) + Fast Green (FG, 2 mM; pressure pulse: 750 ms, 5 PSI; D, compare with C) or to a depolarizing current pulse (+4 nA, 10 s; E) to mimic the depolarization elicited by transient IP3 injection (compare with C). No effect on the ongoing membrane potential oscillation was observed with either protocol, thereby excluding contributions from nonspecific effects of injections (e.g., mechanical cell swelling) or the ability of direct, transient membrane depolarizations to induce phase shifts in oscillation.

The involvement of this organelle-based pacemaker mechanism in the learning-induced plasticity was then further assessed in Non-contingent, Contingent, and Contingent+IP3-injected B63 preparations under chemical synapse blockade, as described earlier. Comparisons were performed during a 10 min test period that began 20 min after the onset of low Ca + Co perfusion, and during which time, all recorded B63 neurons were hyperpolarized to −80 mV by tonic intracellular injection of negative current to maintain their basal membrane potentials at the same resting level and to decrease the likelihood of plateau potential genesis. Recordings were performed after equivalent post-training delay intervals across the groups (Non-contingent, 6.0 hr; Contingent, 5.9 hr; Contingent+IP3, 6.1 hr; H = 1.695, p = 0.689).

Spectral and wavelet analyses showed that Contingent training significantly increased the frequency of B63’s spontaneous oscillation relative to that expressed after Non-contingent training (Figure 7A,B, Figure 8A: H = 16.239, p < 0.001; Contingent vs. Non-contingent, q = 2.055, p = 0.04). This plasticity depended on a change to an organelle-derived pacemaker mechanism, as it was not observed in Contingent+IP3-injected B63 neurons (vs. Contingent, q = 4.029, p < 0.001; vs. Non-contingent, q = 2.157, p = 0.04). In contrast, the oscillation amplitude was not significantly altered by learning (Figure 7A-C; Figure 8B: H = 7.691, p = 0.016; Contingent vs. Non-contingent: q = 0.243, p = 0.808), but again was strongly reduced in the Contingent+IP3 group (vs. Contingent: q = 2.518, p = 0.028; vs. Non-contingent: q = 2.352, p = 0.028). These findings under conditions of functional isolation therefore indicate that the learning-induced increase in the frequency of buccal motor output arises from a sustained acceleration of the endogenous voltage oscillation generated by an intracellular calcium dynamic that is specific to the B63 neurons.

**Figure 7.**
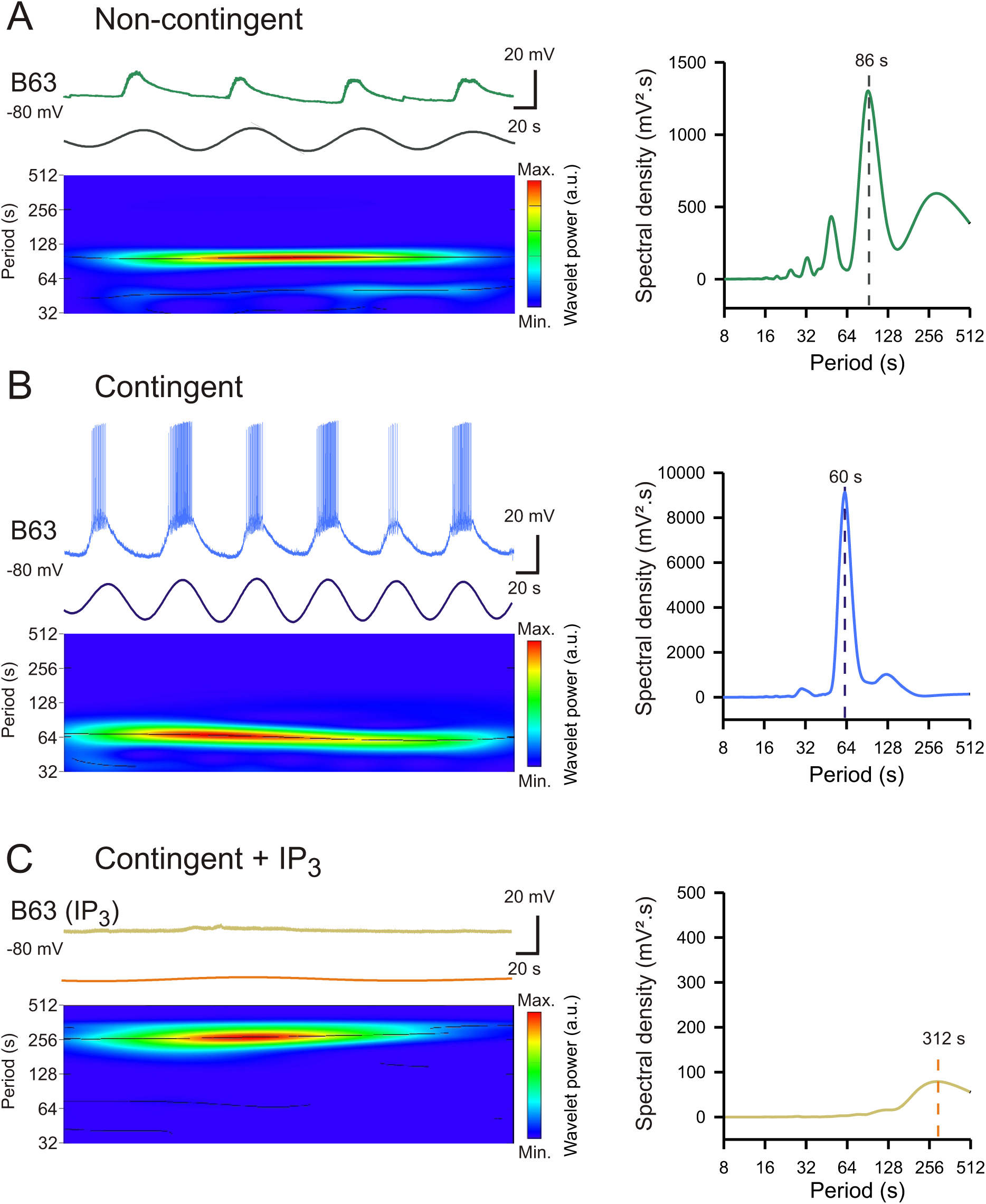
Learning-induced acceleration of B63’s spontaneous, low-amplitude voltage oscillation and the cell’s increased excitability under chemical synapse blockade. (**A-C**) Spontaneous voltage oscillations (left, upper traces) in different B63 neurons from Non-contingent (A) and Contingent (B) preparations, and a further Contingent preparation after injection of IP3 (C), and with synapses blocked with ‘low Ca^2+^ + Co^2+^‘. The 3 neurons were additionally held hyperpolarized to −80 mV (from the hyperpolarized trough between oscillation cycles) by continuous negative current injection to prevent plateau potential production. Colored panels at left: Wavelet analyses of each cell’s voltage fluctuations and reconstructed waveforms (colored sinusoids) indicated by the single dominant period (dashed lines) in the corresponding spectral density plots at right. The intrinsic, low-amplitude oscillations in B63 occurred at a higher frequency in the Contingent preparation (B) than in either the Non-contingent (A) or the Contingent+IP3 (C) preparation. Moreover, although held hyperpolarized, the Contingent B63 cell’s oscillation continued to elicit bursts of full size action potentials, but not the B63s in the other two types of preparation.

**Figure 8.**
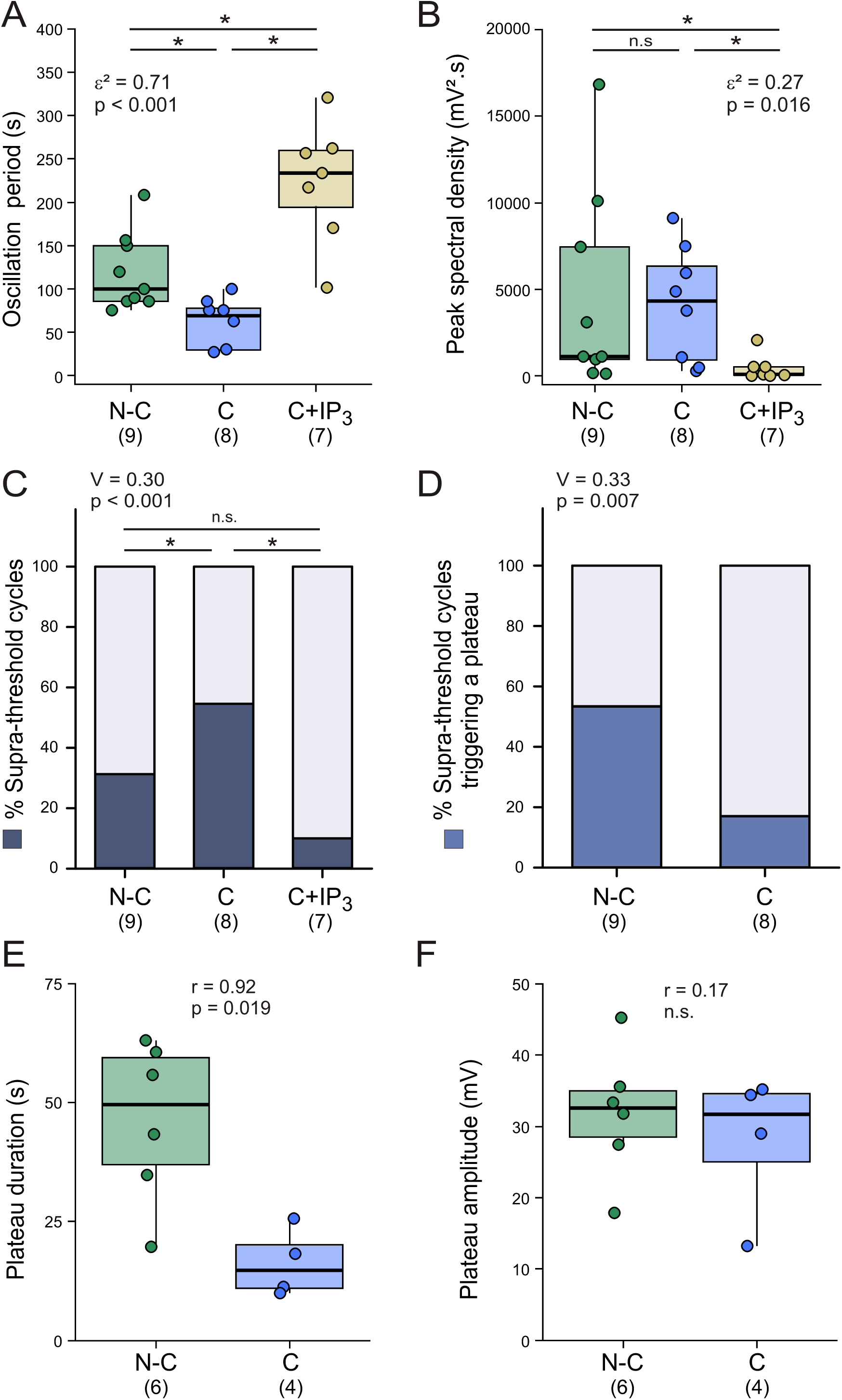
Quantification of the learning-induced plasticity in B63’s spontaneous membrane behavior in the absence of functional chemical synapses. (**A,B**) Between-group comparisons of the frequency (A) and amplitude (B) of spontaneous low-amplitude voltage oscillations in B63 neurons recorded under low Ca^2+^ + Co^2+^ solution and hyperpolarized to −80 mV by continuous current injection. Prior contingent training *in vivo* (middle boxes), compared with non-contingent training (left boxes), significantly decreased B63’s oscillation period without significantly (n.s.) changing its amplitude. In contrast, both the frequency and amplitude of membrane potential fluctuations were significantly decreased in Contingent preparations in which B63 neurons had been iontophoretically injected with IP3 (right boxes). Note that cell impalement was lost before sufficient recording in 1/9 C and 2/9 C+IP3 preparations. (**C,D**) Comparisons of B63’s membrane excitability in the same preparation groups as in A,B. The proportions of suprathreshold oscillation cycles (C, blue shading), and of these that triggered a plateau potential (D; blue shading), differed significantly between groups: compared to the Non-contingent group (left boxes), Contingent training increased the proportion of B63 oscillation cycles that elicited action potentials (C; middle box), although the likelihood of plateau potential production was decreased (D; right box). Following injection of IP3, Contingent B63 neurons displayed markedly reduced impulse production (C) and none of this group produced plateau potentials (thus absent from D). **(E,F)** Group comparisons of B63 plateau potential durations and amplitudes. Contingent training, compared with Non-contingent training, significantly reduced the duration of B63 plateaus when expressed (E) but without altering their amplitude (F).

### Learning induces an associated increase in B63’s membrane excitability

After chemical synapse blockade and during tonic hyperpolarization of the B63 neurons to −80 mV, the likelihood that individual oscillation cycles triggered bursts of full-size action potentials, with or without accompanying large amplitude plateau potentials, differed significantly between the groups (Figure 7A-C; Figure 8C, p < 0.001). Despite the absence of significant changes in oscillation amplitude, the proportion of cycles that were supra-threshold for impulse firing was significantly higher in the Contingent group than in either the other two groups (Contingent vs. Non-contingent: p = 0.020; vs. Contingent+IP3: p = 0.002). No significant difference was found between the Non-contingent vs. Contingent+IP3 groups (p = 0.124).

This learning-dependent decrease in the threshold for spiking activity differentially affected the generation of action and plateau potentials. In the Contingent group compared with the Non-contingent group, whereas the proportion of oscillation cycles producing bursts of full-size action potentials increased, the percentage of cycles additionally triggering plateau potentials significantly decreased (Figure 8D, Contingent vs. Non-contingent: p = 0.007). In Contingent preparations with IP3-injected B63 neurons, the residual low amplitude voltage oscillation rarely elicited action potentials and failed completely to trigger plateau potentials. Moreover, when plateaus were produced in the Contingent group, their durations, but not their amplitudes, were significantly reduced compared with the Non-contingent group (Figure 8E,F; duration: W = 1, p = 0.019; amplitude: W = 10, p = 0.762). Therefore, operant conditioning increases the intrinsic excitability of B63 neurons by promoting impulse firing while reducing the expression of prolonged plateau potentials, indicating that learning leads to an independent modulation of the voltage-dependent channels responsible for these active membrane properties.

These learning-induced changes in B63 neuron excitability could reflect a direct, long-lasting modulation of the plasma membrane channels themselves, or indirectly, from a change in the kinetics of intracellular calcium fluxes underlying the spontaneous oscillation, which in turn affects ion channels in B63’s membrane. To distinguish between these possibilities, the excitability of B63 after *in vivo* training was further tested using experimental intracellular injections of depolarizing current steps applied between the depolarized peaks of two consecutive spontaneous oscillation cycles, during the phase when the resting membrane potential had repolarized to the −80 mV holding potential. In the same groups of preparations analyzed in Figure 8, thresholds for eliciting action potentials and plateau potentials were determined using progressively increasing current intensities (up to 15 nA) with successive pulses of 1 s or 30 s durations (Figure 9A,B).

**Figure 9.**
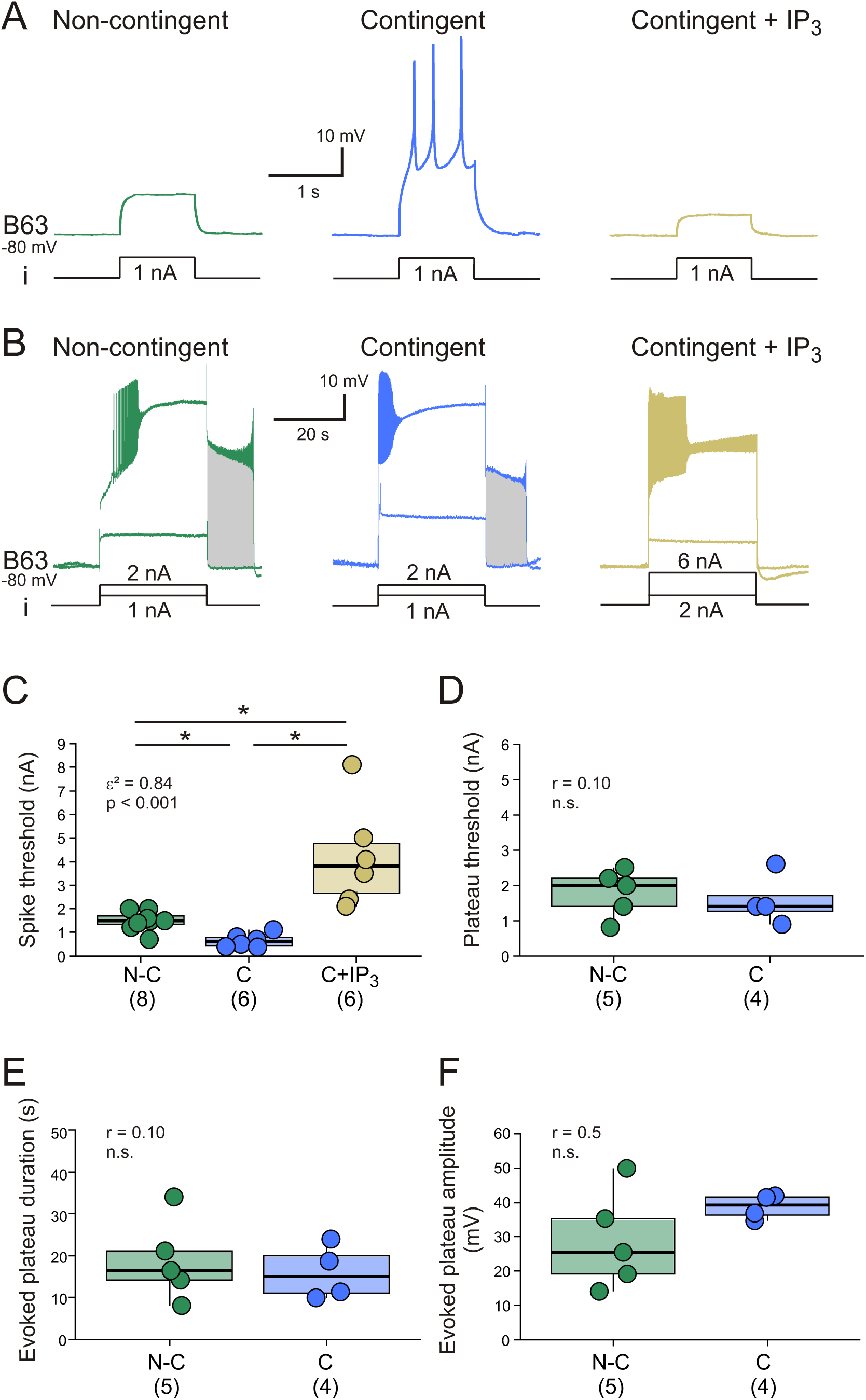
Learning-induced plasticity in B63’s membrane properties assessed by depolarizing current injection. (**A,B**) Determination of activation thresholds for action potentials (A) and plateau potentials (B) using depolarizing current pulses of increasing intensity injected into B63 neurons under chemical synapse blockade. All current injections were performed in the hyperpolarized troughs between successive spontaneous oscillation cycles while the cell’s membrane potential was held at −80 mV by continuous intracellular injection of hyperpolarizing current. Under these conditions, brief depolarizing pulses (i, 1 s) of less current intensity triggered action potentials in B63 of a Contingent preparation compared with either a Non contingent or Contingent + IP3 preparations (A). In contrast, depolarizing pulses (i, 30 s) of similar intensities were required to trigger plateau potentials that outlasted the current pulse before a repolarization to baseline membrane potential (B, shaded areas) in both Contingent and Non-contingent preparations. No prolonged plateau potentials could be elicited with depolarizing current pulses up to 15 nA in the B63 neuron injected with IP3 after Contingent training (B, right). (**C,D**) Between group comparisons of thresholds for action potentials (C) and plateau potentials (D). Contingent learning significantly decreased impulse thresholds (C, middle box) compared with Non-contingent training (C, left box). Such an increased excitability was absent in Contingent IP3-injected neurons, which instead exhibited significantly higher action potential thresholds (C, right box). In contrast, no Contingent training-related change in plateau activation thresholds was evident (D), and no plateaus were produced by IP3-filled neurons. Note that cell impalement was lost before or during testing in 1/9 N-C and 3/9 C and 3/9 C+IP3 preparations. Moreover, B63 neurons from 3/8 N-C, 2/6 C, and 6/6 C+IP3 preparations failed to produce plateau potentials in response to current pulse intensities up to 15 nA. **(E,F)** Group comparisons of the duration (E) and amplitude (F) of current pulse-evoked plateau potentials. Under the experimental conditions described for A,B, no significant differences in the post-pulse duration of evoked plateau potentials and their amplitudes were found between the Contingent and Non-contingent groups.

Full-size action potentials were consistently elicited by shorter and lower-intensity pulses than plateau potentials, regardless of the prior training protocol. Moreover, less current intensity was required to reach B63’s impulse threshold in the Contingent preparations compared with either Non-contingent or Contingent+IP3 preparations, whereas more current was needed in IP3-loaded B63 cells than in neurons in either non-injected group. These results, which are consistent with the altered responsiveness to spontaneous depolarizations (Figure 7, 8C), thus indicate that B63’s membrane excitability was enhanced by operant conditioning and down-regulated by elevated IP3 (Figure 9A, C; H = 15.888, p < 0.001; Contingent vs. Non-contingent: q = 1.965, p = 0.049; vs. Contingent+IP3: q = 3.981, p < 0.001; Non-contingent vs. Contingent+IP3: q = 2.291, p = 0.033).

In contrast, learning had no effect on the threshold (Figure 9B,D; W = 9, p = 0.901) or on the amplitude and duration of current pulse-evoked plateau potentials, as indicated by the delay to spontaneous repolarization following pulse termination (Figure 9E,F; duration, W = 9, p = 0.905; amplitude, W = 15, p = 0.286), whereas IP3 injection abolished the expression of sustained plateaus that outlasted current injection. The absence of a learning-related change in current-evoked plateau durations therefore contrasted with the significant plateau shortening occurring during spontaneous oscillations (see Figure 8E). This finding, together with the effect of IP3, are therefore consistent with the earlier conclusion that any learning-related plasticity in B63’s plateau generating capability may depend on changes to intracellular calcium dynamics that regulate the membrane channels responsible for plateau termination (Bédécarrats et al., 2023; see Discussion).

## Discussion

This study investigated the contribution of slow, spontaneous voltage oscillations in an homologous pair of decisional interneurons to the expression of a compulsive-like motor behavior after associative reward learning. Operant conditioning transforms infrequent and irregular motor activity responsible for *Aplysia’s* food-seeking movements into frequent and regularly recurring motor pattern cycles, accompanied by accelerated membrane potential oscillations and an increased excitability of these two key constituents of the buccal CPG network. This plasticity, which is critically dependent on changes to an intracellular calcium oscillation arising from organelle calcium release and reuptake in these specific neurons and the activation of certain plasma membrane channels, is governed by the second-messenger IP3. Our findings therefore indicate that organelle calcium-driven voltage oscillations and membrane excitability provide the basis for cellular memory traces underlying compulsive-like motor behavior induced by operant conditioning.

### Role of spontaneous voltage oscillations in learning and memory

Spontaneous voltage oscillations in neuronal circuitry with periods of seconds to minutes are widely involved in olfactory processing in vertebrates and invertebrates, and can be modulated by appetitive and aversive classical conditioning (Friedman and Strowbridge, 2003; Gelperin and Tank, 1990; Kay, 2015; Rosay et al., 2001; Sekiguchi et al., 2010). In these learning paradigms, pairing odors with unconditioned stimuli modifies the frequency, amplitude or spatial propagation of the oscillatory responses to conditioned odors alone, thereby leading to a subsequent modulation of olfactory-driven behavior (Inoue et al., 2006; Kimura et al., 1998; Samarova and Balaban, 2009; Watanabe et al., 2008).

Spontaneous voltage and associated calcium oscillations are also involved in the consolidation of long-term memories. In humans and rodents, the dynamics of slow electrical rhythms generated by cortical networks that emerge when the flow of sensory inputs is decreased can be modulated by learning (Huber et al., 2004; Kattler et al., 1994; Klinzing et al., 2019; Miyamoto et al., 2017; Mölle et al., 2004). Experimental stimulation or suppression of this oscillatory activity can facilitate or disrupt long-term memory formation and retrieval (Gulati et al., 2017; Marshall et al., 2006; Ngo et al., 2013). Similar causal relationships between network oscillatory activity and the formation or consolidation of long-term memory have also been established in invertebrates. For example, spontaneous slow calcium oscillations are expressed by Kenyon cells located within *Drosophila*’s mushroom bodies and in projection neurons to this memory center. Learning paradigms or alterations in the expression of the *amnesiac* gene, which is required for memory consolidation, modulate the frequency or amplitude of these oscillations in correlation with persistent changes in long-term memory storage (Plaçais et al., 2012; Rosay et al., 2001). Thus, long-lasting changes to spontaneous, slow oscillatory processes in sensory and/or other circuits that are upstream to motor centers can be involved in the transformation of past experiences into long-term memories that guide future behavior.

Our present findings show that slow, spontaneous voltage oscillations in identified constituents of an *Aplysia* CPG circuit that actually drives behavior, are not only necessary for generating motor output but also contribute to the memory formation produced by operant conditioning. This rhythmic pacemaker signal, which originates uniquely in the B63 neurons of the buccal network, arises from a cyclic release/reuptake of calcium from intracellular stores and is converted into membrane potential oscillations through the activation of calcium-activated conductances (Bédécarrats et al., 2021, 2023). The pacemaker oscillation, which is assumed to arise remotely from B63’s soma in electrotonically connected neuropilar regions of the cell (Bédécarrats et al, 2021), then spreads through gap junctions to electrically- and metabolically-coupled buccal network partners to produce coordinated membrane potential fluctuations throughout this neuronal subpopulation and a resultant initiation of cycles of network-wide activity. In ganglia from unconditioned *Aplysia,* B63’s voltage oscillation expresses cycle-to-cycle variations in amplitude that fluctuate around the thresholds for action and plateau potential generation, resulting in sporadic and infrequent network activation and motor output production. In contrast, after operant conditioning, the frequency of B63’s oscillation is persistently increased in association with an increase in membrane excitability in the B63 neurons and their electrically-coupled network partners (Costa et al., 2020; Sieling et al., 2014). Consequently, the more reliably and frequently repeating impulse bursts occurring synchronously throughout this subcircuit lead to the triggering of accelerated and regularly recurring BMP cycles.

Although this learning-induced plasticity is expressed by still active isolated buccal ganglia, in our *in vitro* experiments to date, monotonic electrical stimulation of buccal nerves carrying sensory axons to the ganglia was systematically used to replicate the inciting food stimulation applied to the intact animal during training (Nargeot et al., 2007, 2009). This raises the possibility that at least some of the alterations in buccal network operation are mediated indirectly by changes at the central synapses of these input pathways, and that these memory traces then impact on B63 and the wider CPG circuit during subsequent pathway activation *in vitro*. Indeed, there is a vast literature on the plasticity of sensory to motoneuron and central circuit synapses resulting from associative and non-associative learning in a wide variety of animal model systems, and *Aplysia* in particular (Tam et al., 2020; Hurwitz et al., 2024; for review see Kandel, 2001; Byrne and Hawkins, 2015; Hawkins and Byrne, 2015). However, our finding that the buccal network can continue to function spontaneously and express learning-related changes in the complete absence of sensory nerve stimulation indicated that any such influences do not play a major role, but rather, the memory traces for appetitive operant conditioning are encoded within the CPG circuit and specifically the B63 neurons (see also Costa et al., 2022).

### Learning-induced plasticity of B63’s intracellular calcium oscillation

Previous evidence that the second messenger IP3 plays a critical role in generating B63’s intracellular calcium oscillation derived from the injection of an IP3 receptor antagonistic, heparin, to block organelle calcium release or the reticulum calcium pump inhibitor, CPA (Bédécarrats et al., 2021), both of which suppressed the neuron’s low amplitude voltage oscillation. In the present study, we used direct injections of IP3 itself, with pulsed or continuous applications to assess its effects on B63’s bioelectrical behavior and the plasticity resulting from operant learning. The two approaches provided further insights into the neuron’s endogenous pacemaker mechanism and its modification by learning. Firstly, individual pressure-injected pulses of IP3 caused a transient membrane depolarization and a subsequent resetting of B63’ ongoing voltage oscillation, which is a defining feature of biological oscillators in general (Ayers and Selverston, 1977; Fenk et al., 2024; Glass and Mackey, 1988; Winfree, 1977). The lack of equivalent effects of transient depolarizing current injection indicated that this oscillatory property does not rely on voltage-dependent conductances in the plasma membrane, which is in accordance with the earlier finding that the frequency of B63’s oscillation remains unaltered by continuous experimental depolarization or hyperpolarization (Bédécarrats et al, 2021). Rather, the responses we observed with IP3 injection are consistent with the canonical classification of the so-called type 2 category of IP3-dependent oscillator whereby the calcium oscillation, accompanied by oscillations in intracellular levels of IP3 itself, can occur only within an appropriate range of IP3 concentration (Harootunian et al., 1991; Sneyd et al., 2006). On this basis, and to which our B63 data also comply, a transient increase in IP3 concentration via an external pulse, elicits a rise in intracellular calcium followed by a suppression of the oscillation and a resultant phase shift when the oscillation reappears as the IP3 level has sufficiently decreased.

Our second approach in using exogenous IP3, which was employed in most experiments reported here, was to apply the second messenger via continuous iontophoretic injection. Such constant intracellular loading, which presumably clamped IP3 at a near maximal concentration for oscillator function, consistently caused a drastic reduction in both the amplitude and frequency of B63’s oscillation, likely due to a sustained near-inactivation of the cell’s organelle IP3 receptors by the elevated levels of store released calcium (Hajnóczky et al., 1995; Wakui et al., 1989). Significantly, moreover, the residual slow oscillation that persisted in IP3-injected B63 neurons no longer expressed the cycle frequency increase seen in their non-injected counterparts after operant conditioning, further indicating that long-lasting changes in the dynamics of cytosolic, IP3 receptor-controlled calcium handling make a crucial contribution to the actual learning process. Whether or not the memory trace is transiently suppressed or permanently removed by IP3’s presence in B63, which would require experimental retesting after the neuromodulator’s post-injection removal from the cell’s cytoplasm, remains to be determined.

Learning has been found to regulate second-messenger-dependent oscillations, including calcium oscillations, in other neuronal systems of both vertebrates and invertebrates (Berridge, 2014; Deng et al., 2025; Plaçais et al., 2012; Rosay et al., 2001). These oscillations are generally thought to arise from calcium influx through voltage-gated membrane channels, a regulation of intracellular calcium concentration by organelles, and feedback control of plasma membrane conductances (Berridge, 2014; Crunelli et al., 2005; Yu et al., 2004). Moreover, the frequency of such oscillatory activity can be modulated by neurotransmitters that change membrane channel properties, second-messenger concentrations, and organelle receptor kinetics (Canavier et al., 1991; Chevalier et al., 2016; Dalla Porta et al., 2025; Dupont et al., 2011; Sneyd et al., 2006, 2017; Zhao et al., 2010). Similarly, in *Aplysia*, the modulation of B63’s organelle-derived calcium oscillation by operant conditioning could involve monoamines released by food-activated reward pathways, which act on intracellular calcium fluxes and/or plasma membrane properties (Brembs et al., 2002; Butera et al., 1995; Kadiri et al., 2011; Martínez-Rubio et al., 2009; Nargeot et al., 1999b).

### Multiple sites for memory storage in rhythmogenic networks

Several forms of learning define the activity patterns of rhythmic networks, including *Aplysia’s* buccal circuit, by modifying chemical and electrical synapses as well as the membrane properties of circuit neurons (Braun and Lukowiak, 2011; Costa et al., 2020; Hurwitz et al., 2024; Kemenes et al., 2006; Klinzing et al., 2019; Li et al., 2024; Marra et al., 2010; McComb et al., 2005; Miyamoto et al., 2017b; Sieling et al., 2014; Spencer et al., 1999; Staras et al., 2003; Tam et al., 2020; Yoshida and Hayashi, 2004). Operant-reward conditioning of *Aplysia’s* feeding behavior has been shown to change chemical synaptic strength and the membrane excitability amongst certain decisional buccal network neurons (B51, B4/B5), whose activity generates key features of the motor program for the behavior that was contingently rewarded during prior learning. This combined learning-induced plasticity thereby contributes to the selection of buccal network output by biasing circuit operation towards the production of food ingestion movements rather than unrewarded actions such as egestion behavior (Brembs et al., 2002; Costa et al., 2022, 2020; Momohara et al., 2022; Mozzachiodi et al., 2008; Nargeot et al., 1999a). Moreover, operant learning modifies the gap junction coupling amongst the buccal network subset (bilateral B63, B30, B65) that is decisional for triggering the initial protraction phase of each radula BMP. Although electrical coupling does not contribute to actual rhythmogenesis, which is strictly endogenous to the B63 cells, a learning-induced increase in the strength of coupling between these neurons, in association with an increased excitability, synchronizes their enhanced bursting activity, thereby increasing the probability of frequent and regular BMP initiation (Costa et al., 2020; Nargeot et al., 2009; Sieling et al., 2014).

Our present study further establishes that the membrane excitability of the B63 neurons is altered by learning. The threshold for action potential generation, whether triggered by B63’s spontaneous oscillation or by depolarizing current injection, was found to be lowered in buccal ganglia from animals subjected to operant conditioning. Conversely, the threshold for plateau potential initiation appeared to remain unaffected. However, the duration of plateaus evoked by B63’s endogenous voltage oscillation was significantly decreased by learning, although not the duration of sustained plateaus that had been triggered experimentally by transient depolarizing current injection. It is feasible, therefore, that the activation thresholds for conductances responsible for active plateau termination are modulated by learning. Likely candidates are voltage-dependent, calcium-activated potassium channels, which are known to contribute to plateau termination in both invertebrate (Golowasch and Marder, 1992; Neveu et al., 2023) and vertebrate neurons (Cai et al., 2004; el Manira et al., 1994; Russo and Hounsgaard, 1996), including *Aplysia’s* B63 cells (Bédécarrats et al., 2023). Moreover, a learning-related modulation of this channel type has been widely reported (Mpari et al., 2010; Pham et al., 2023; Titley et al., 2020; Yamoah and Crow, 1995). In *Aplysia’* B63 cells, a learning-induced increase in levels of cytosolic calcium during oscillation and/or long-lasting decrease in the activation threshold of KCa channels themselves could explain the reliable plateau termination soon after each peak of spontaneous oscillation. The prevention of plateau prolongation would in turn promote the reoccurrence of a short impulse burst -and a BMP initiating protraction phase -in the next oscillation cycle.

Therefore, in addition to central decision-making processes determining the selection and cycle-by-cycle initiation of buccal motor output for different feeding-related behaviors, learning can modify the neuronal mechanisms that are responsible for autonomously generating these patterns. It must be emphasized that B63’s organelle-derived calcium dynamic serves as an oscillator mechanism underlying the expression of a motivated behavior. Thus, we believe that our findings that this spontaneous intracellular process is modified by operant conditioning provides the first reported example of such a second-messenger-dependent mechanism playing a key role both in the rhythmogenesis of the central network responsible for a behavior and in the latter’s plasticity resulting from learning.

In conclusion, an acceleration of an endogenous calcium dynamic in the decision-making B63 neurons, the resulting membrane voltage oscillation that more reliably generates bursting, and its synchronized propagation through strengthened gap junctions to network partners, constitute multiple facets of the memory arising from operant conditioning. The outcome of these cellular and synaptic changes for *Aplysia’s* motivational search for food is that an otherwise random and impulsive motor act is transformed by operant experience into a compulsive-like expression of this behavior.

## Materials and Methods

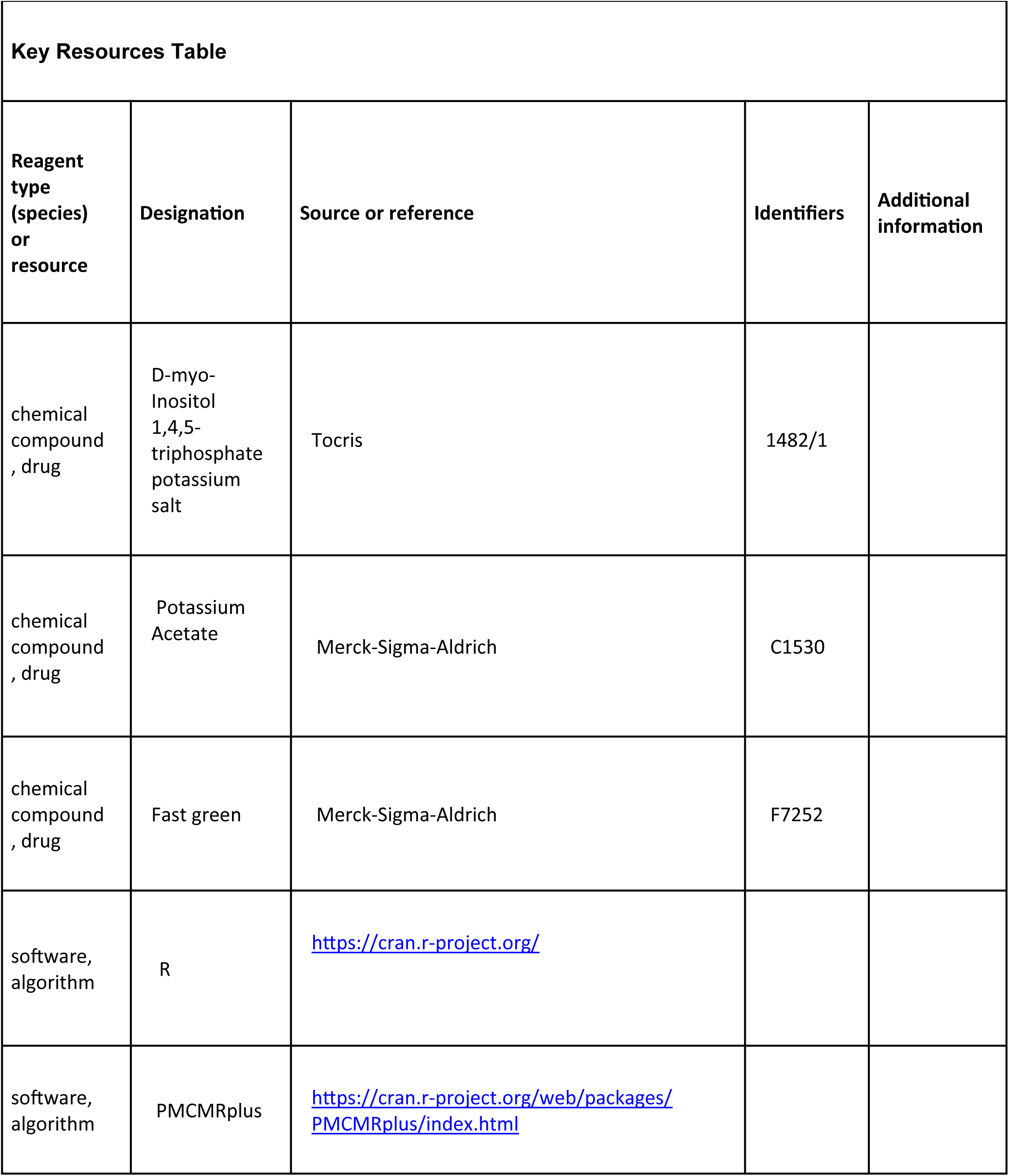
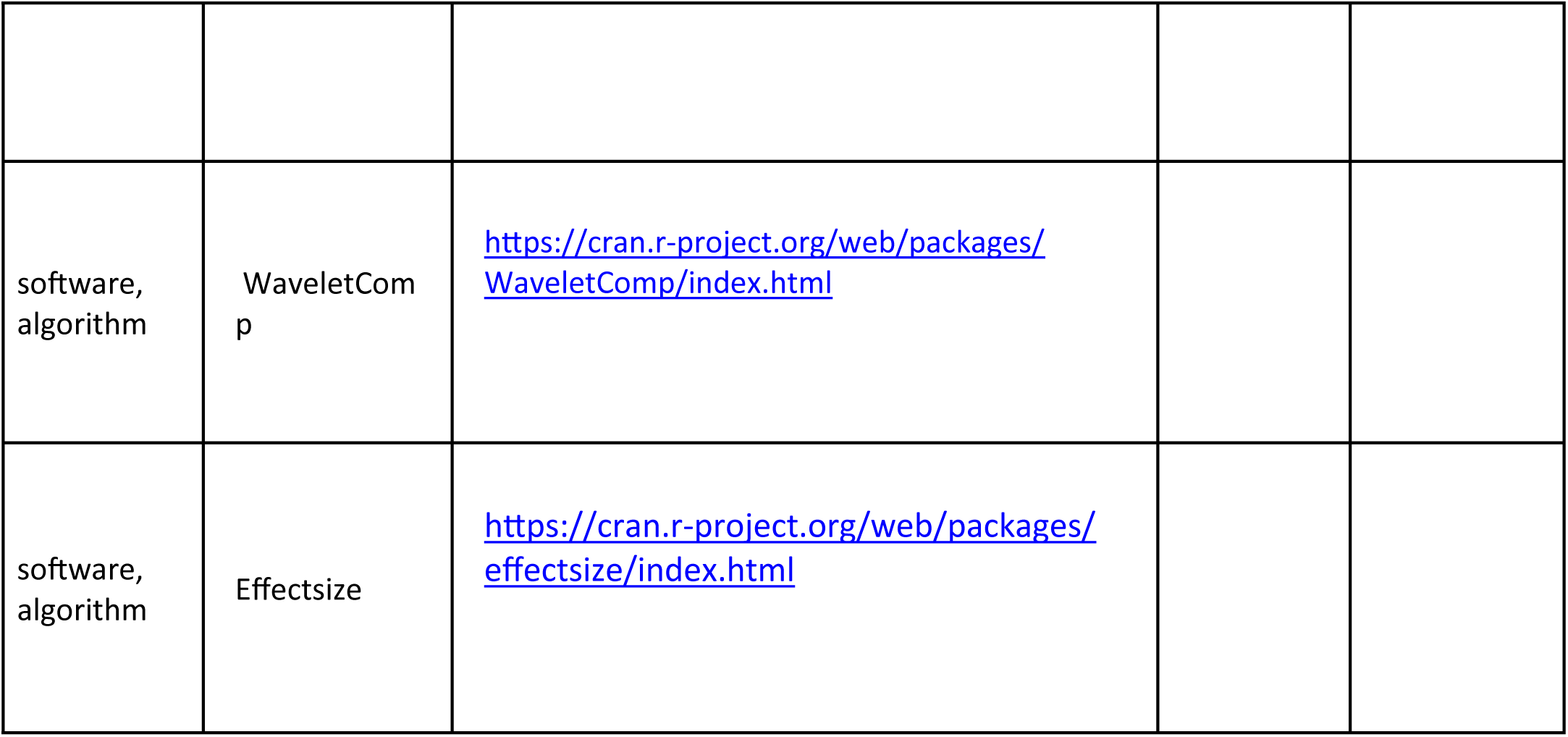

| **Reagent type (species) or resource** | **Designation** | **Source or reference** | **Identifiers** | **Additional information** |
|---|---|---|---|---|
| chemical compound, drug | D-myo-Inositol 1,4,5-triphosphate potassium salt | Tocris | 1482/1 |  |
| chemical compound, drug | Potassium Acetate | Merck-Sigma-Aldrich | C1530 |  |
| chemical compound, drug | Fast green | Merck-Sigma-Aldrich | F7252 |  |
| software, algorithm | R | https://cran.r-project.org/ |  |  |
| software, algorithm | PMCMRplus | https://cran.r-project.org/web/packages/PMCMRplus/index.html |  |  |
| software, algorithm | WaveletComp | https://cran.r-project.org/web/packages/WaveletComp/index.html |  |  |
| software, algorithm | Effectsize | https://cran.r-project.org/web/packages/effectsize/index.html |  |  |

### Animals and behavioral training

All experiments were performed on adult *Aplysia fasciata* (100-200 g, hermaphrodite) collected locally from the Bassin d’Arcachon, France, and housed in tanks containing fresh aerated seawater (∼15°C). Animals were fed with fresh or frozen seaweed (*Ulva lactuca* or *Carradoriella elongata* supplied by the Station Biologique at Roscoff, France). Behavioral training procedures followed those described previously in detail (Nargeot et al., 2007; see also the Results section). Animals were randomly assigned to the different experimental groups. Briefly, feeding behavior was incited by the continuous application of a none-ingestible food stimulus (*Ulva lactuca*) to the animals’ lips. A rewarding food stimulus consisted of 20 µL of seaweed juice prepared by macerating 0.4 g of dried *U. lactuca* in 10 ml of artificial sea water (ASW) for 1 hr. This second stimulus was delivered via a pipette, either (1) contingent upon each occurrence of a radula biting cycle (‘Contingent’ groups of animals), or (2) independently of the expression of this autonomous behavior, but with the same number of food reward deliveries as each paired Contingent animal and at regular time intervals (‘Non contingent’ control groups).

### Salines, electrophysiology, and intracellular drug injections

Saline compositions, intra- and extra-cellular electrophysiological recording and stimulation methods applied to isolated buccal ganglia were as those described in Bédécarrats et al. (2021). Intra-somatic injections were performed with either iontophoretic or pressure injection techniques, using glass micro-electrodes backfilled for ∼5 min with a filtered (0.2 µm) solution of D-myo-inositol 1,4,5 triphosphate potassium (IP3; 10 mM, Tocris), control potassium acetate (10 mM) containing Fast Green (Fast Green, 2 mM; Merck-Sigma-Aldrich), or Fast Green alone (10 mM). For iontophoretic injections, negative current pulses (-2 nA, 500 ms duration, 1 Hz) were delivered for 1 hr using a Grass S88 stimulator. Pressure injections were performed with air pulses of 5-10 PSI, 150-750 ms durations, generated by a Picospritzer II. In this latter procedure, electrode tips were broken mechanically to produce a tip resistance of 10-40 MΩ. Electrodes that did not meet this criterion or failed to eject solution efficiently into the preparation bathing medium (ASW) were replaced before further experimentation. Following soma impalement, data were excluded and electrodes replaced if effective intracellular injection was not confirmed either by visible cell staining (for Fast Green-containing electrodes) and/or by a cellular electrophysiological response.

### Quantification and Statistical analyses

The variability (or regularity) of biting behavior was evaluated by testing for a significant Gabor sinusoidal function fit to the auto correlogram of inter-bite intervals and quantified by the coefficient of variation (CV) of these intervals, as previously described (Nargeot et al., 2007, 2009). In *in vitro* experiments, because of the relatively low occurrence of spontaneous buccal motor pattern (BMP) generation during a 10 min test period, the variability of motor output was quantified by the CV only when at least three BMPs were expressed. The thresholds for action or plateau potential generation in recorded B63 neurons were determined from an initial resting potential held at −80 mV using a two-electrode current-clamp technique, and was defined as the minimum intracellular depolarizing current required to elicit these responses. The duration and amplitude of spontaneous or evoked plateau potentials were quantified as previously described in Bédécarrats et al. (2023). A burst of action potentials was defined as a discrete sequence of action potentials generated at a frequency > 1 Hz and lasting > 2 s.

Signal analyses of intracellularly-recorded fluctuations in membrane potential of B63 neurons were performed using the R-CRAN “Base” (R Core Team, 2023) and “WaveletComp” packages (Roesch and Schmidbauer, 2018) after first smoothing the raw voltage traces with a Spike 2 “Smooth” filter to suppress action potentials. The peak magnitudes of spectral densities were then determined and the waveform(s) corresponding to oscillations detected on the Fast Fourier Transform (FFT) periodograms were reconstructed from wavelet decomposition, as also previously described (Bédécarrats et al., 2021).

Statistical comparisons were conducted with the R-CRAN “Base”, “PMCMRplus” (Pohlert, 2024) and “effectsize” (Ben-Shachar et al., 2020) packages. For continuous variables, between-group comparisons were carried out using two-tailed Mann-Whitney tests (for two independent groups; W statistic) or Kruskal-Wallis tests (for >2 independent groups, H statistic), followed by Dunn’s *post-hoc* multiple pairwise comparisons with Benjamini-Hochberg correction of p-values (q statistic). Between-group comparisons of proportions were performed using Fisher’s exact tests. When more than two groups were compared, Fisher’s exact tests were followed by multiple pairwise comparisons with Benjamini-Hochberg correction. The strength of a relationship between independent and dependent variables was quantified by effect sizes. These were computed as the rank biserial correlation (r) for the Mann-Whitney test, the rank epsilon squared (ε²) for the Kruskal-Wallis test, and Cramer’V (V) for the Fisher’s exact test, with values ranging from 0 to 1, and with higher values indicating stronger relationships.

Box-plot representations in all figures display median values (bold horizontal lines), along with the first and third quartiles (boxes edges; inter-quartile range, IQR). Outliers, which were not excluded from our analyses, correspond to data points exceeding 1.5 x IQR from the upper or lower quartile (whiskers). Sample sizes (n) are indicated in parentheses below each corresponding plot. Statistical differences were considered significant for p-value ≤ 5% (symbol: *).

## Acknowledgements

This research was supported by a doctoral studentship (to L.P.) from the French ‘Ministère de l’Enseignement Supérieur et de la Recherche’.

## Data availability

The source data files are publicly accessible at : https://entrepot.recherche.data.gouv.fr/dataset.xhtml?persistentId=doi:10.57745/9NCESI

---

**Figure 2-figure supplement 1.**
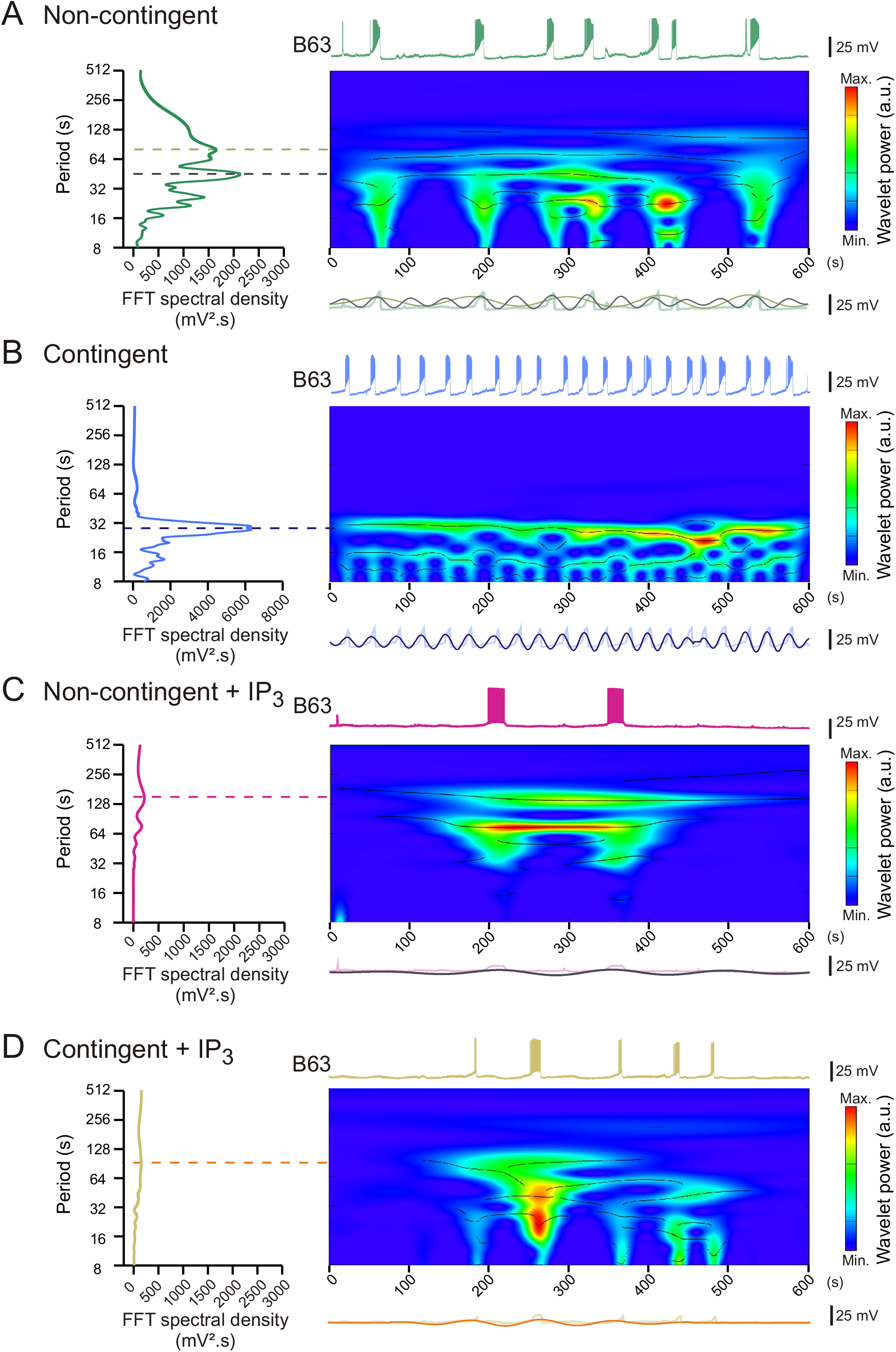
Spectral density and wavelet analyses of B63 membrane potential oscillations during tonic input nerve stimulation. **(A-D)** Top traces at right: representative intracellular recordings from different B63 neurons in Non-contingent (A), Contingent (B) preparations and after injection of IP3 (C,D), as illustrated in Figure 2A1-D1. Colored panels at right: wavelet-based spectral decompositions of B63 membrane potential variations, showing the relative amplitudes of cycles of depolarization over time (color bars; black lines indicate ridges of local maximum power). Left plots: Spectral density analyses (Fast Fourier Transform) used to identify dominant rhythms in the B63s’ voltage fluctuations (dashed lines). Bottom traces at right: reconstructed wavelets from the peak spectral densities (colored sinusoids) superimposed on the corresponding smoothed B63 recording (faded traces).

**Figure 4-figure supplement 1.**
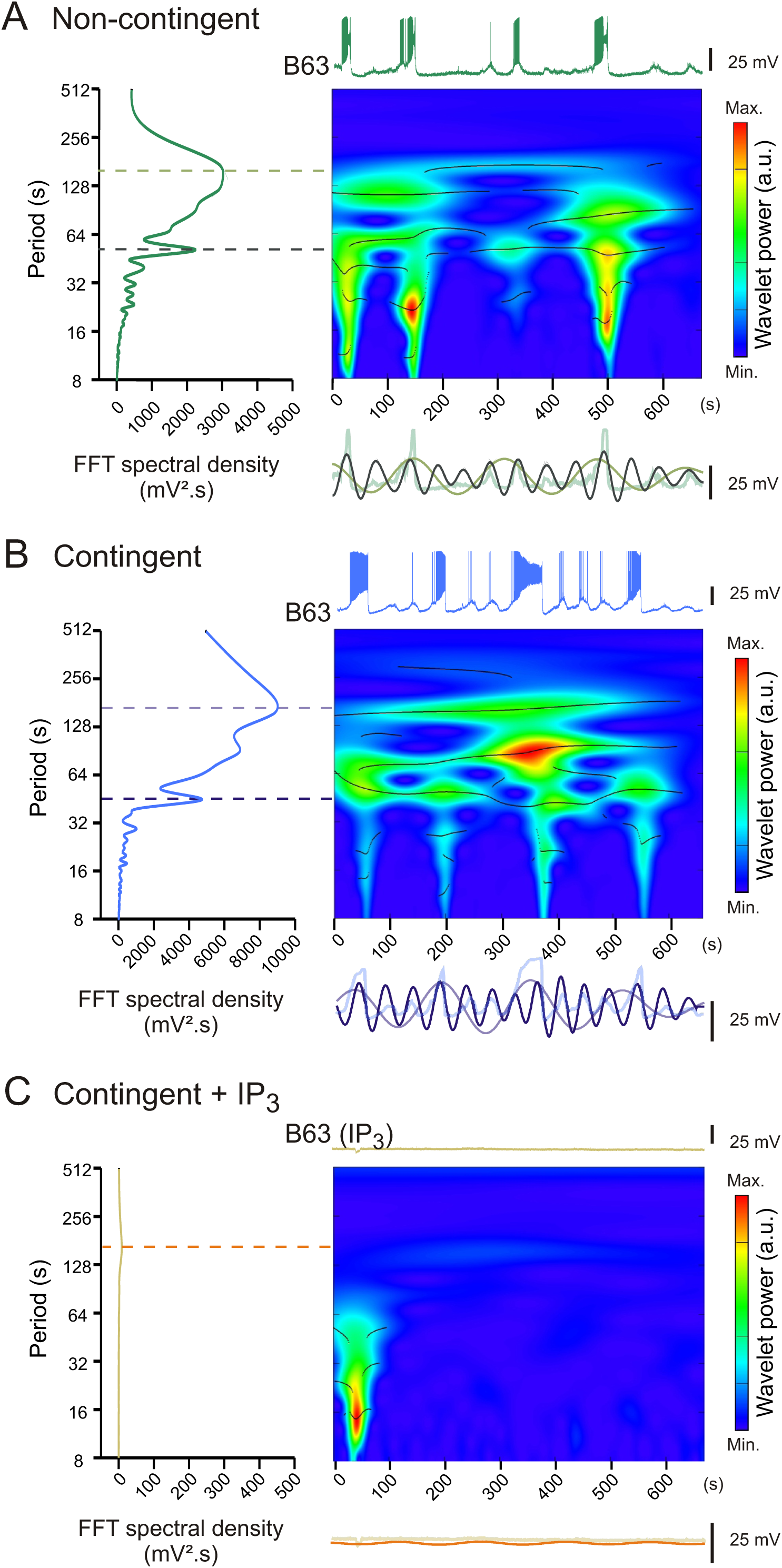
Spectral density and wavelet analyses of spontaneous membrane potential oscillations in B63 neurons. **A-C**) Spectral (periodograms at left) and wavelet (colored panels, center right) analyses of membrane potential fluctuations in different B63 cells (top and bottom traces) as illustrated in Figure 4A-C, respectively (see methodological details in Figure 2-figure supplement 1).

**Figure 6-figure supplement 1.**
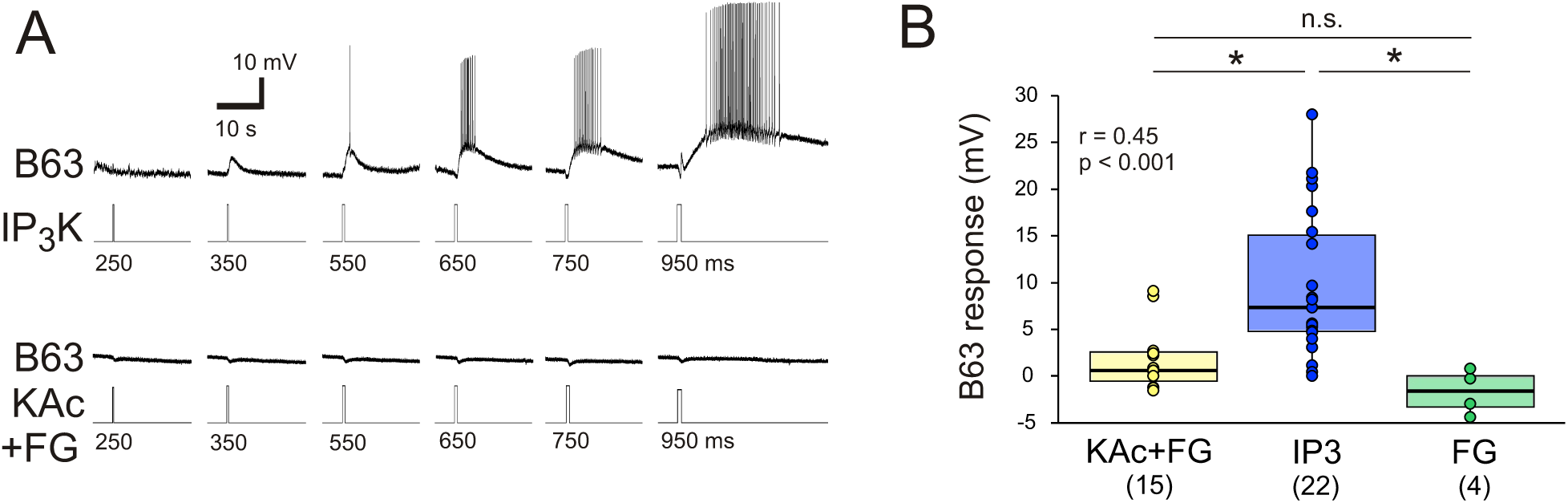
Single oscillation cycles triggered by pulsed pressure injection of IP3 into B63 neurons under chemical synapse blockade. **(A)** Top: Increasing depolarizing responses of the same neuron B63 to successive brief intracellular injections of IP3K (10 mM) delivered with air-pressure pulses of increasing duration (250-950 ms, 5 PSI) under bath-applied ‘low Ca^2+^ + Co^2+^‘ solution to block chemical synapses. Bottom: Control lack of B63 responsiveness in a different preparation to the intracellular injection of potassium acetate (KAc, 10 mM) + Fast Green (FG, 2 mM) with pulses over the same range of durations. **(B)** Comparison of B63 membrane potential responses to intracellular injections of Kac + FG (10 + 2 mM), IP3K (10 mM), or FG alone (10 mM) using 750 ms, 5 PSI air-pressure pulses in independent groups of preparations.

**Figure 6-figure supplement 2.**
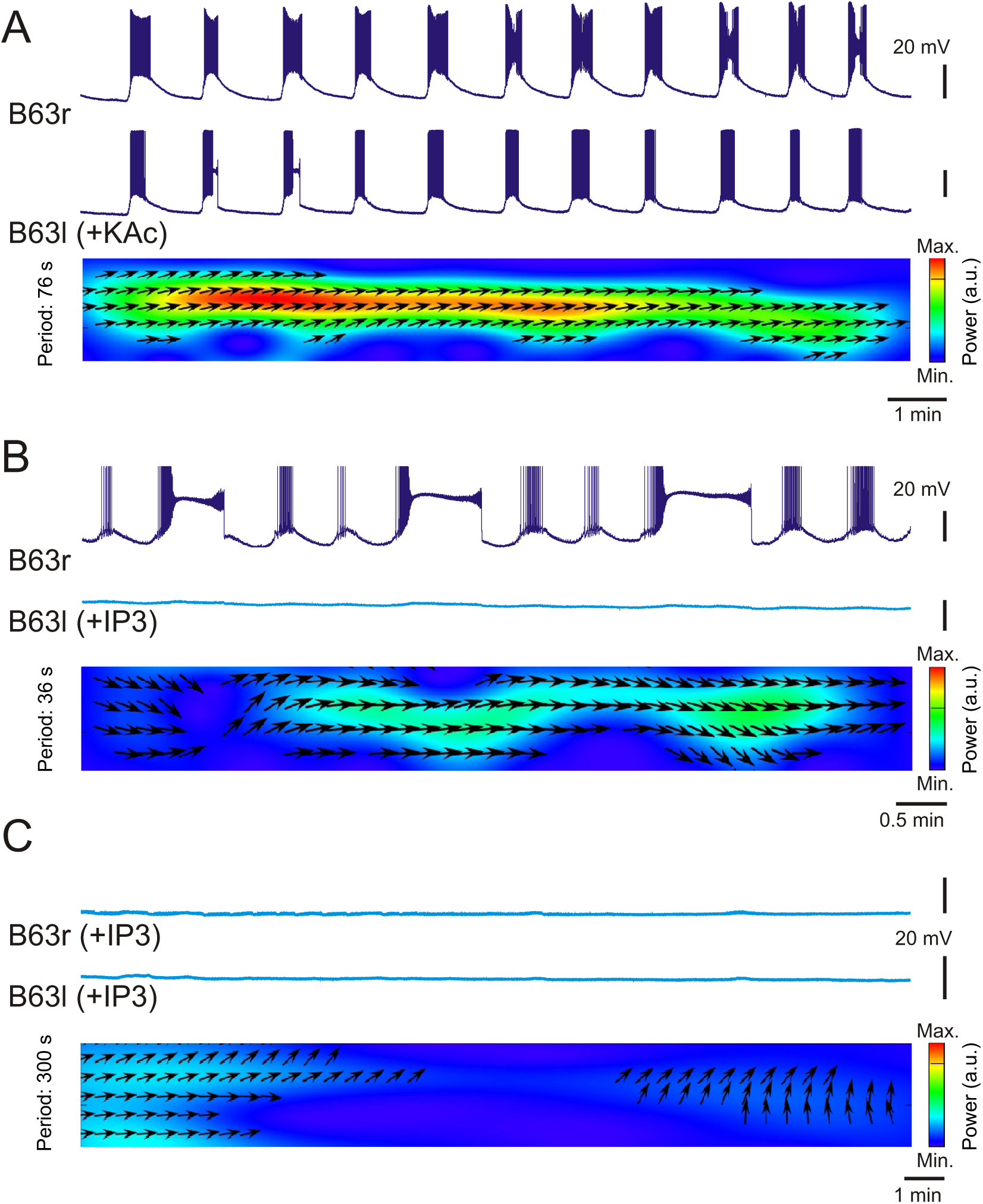
Sustained suppression of B63’s low-amplitude voltage oscillations following iontophoretic IP3 injection. **(A)** Simultaneous intracellular recordings from the two bilateral B63 neurons in a preparation under Low Ca^2+^ + Co^2+^ saline to block chemical synapses, and after a control 1 hr iontophoretic injection (-1.5 nA, 500 ms pulse at 1 Hz) of potassium acetate (KAc, 10 mM) + Fast Green (2 mM) into the left B63 neuron (B63l, bottom trace). Colored panel: Cross-wavelet analysis of the voltage fluctuations in both neurons. Arrows indicate the instantaneous phase relationship (direction towards right: in phase). **(B,C)** Simultaneous recordings from the bilateral B63 neurons in two further different preparations after 1 hr unilateral (B), or bilateral (C) iontophoretic injection of IP3K (10 mM). Colored panels: Cross-wavelet analyses of the voltage fluctuations in the paired recordings (arrow direction towards right: in phase; upwards: phase delay). The unilateral IP3 injection suppressed spontaneous oscillations in the IP3 loaded B63 cell but not the ongoing oscillation and plateau driven bursting in its contralateral, non-injected partner (B), whereas the bilateral IP3 injection completely suppressed the voltage fluctuations of both cells (C).

